# Gut microbiome-linked metabolites in the pathobiology of depression and anxiety - a role for bile acids

**DOI:** 10.1101/2022.04.04.485514

**Authors:** Siamak MahmoudianDehkordi, Sudeepa Bhattacharyya, Christopher R Brydges, Wei Jia, Oliver Fiehn, A John Rush, Boadie W Dunlop, Rima Kaddurah-Daouk, the Mood Disorders Precision Medicine Consortium

## Abstract

**Background:** The gut microbiome may play a role in the pathogenesis of neuropsychiatric diseases including major depressive disorder (MDD). Bile acids (BAs) are steroid acids that are synthesized in the liver from cholesterol and further processed by gut-bacterial enzymes, thus requiring both human and gut microbiome enzymatic processes in their metabolism. BAs participate in a range of important host functions such as lipid transport and metabolism, cellular signaling and regulation of energy homeostasis. BAs have recently been implicated in the pathophysiology of Alzheimer’s and several other neuropsychiatric diseases, but the biochemical underpinnings of these gut microbiome-linked metabolites in the pathophysiology of depression and anxiety remains largely unknown.

**Method:** Using targeted metabolomics, we profiled primary and secondary BAs in the baseline serum samples of 208 untreated outpatients with MDD. We assessed the relationship of BA concentrations and the severity of depressive and anxiety symptoms as defined by the 17-item Hamilton Depression Rating Scale (HRSD_17_) and the 14-item Hamilton Anxiety Rating Scale (HRSA-Total), respectively. We also evaluated whether the baseline metabolic profile of BA informs about treatment outcomes.

**Results:** The concentration of the primary BA chenodeoxycholic acid (CDCA) was significantly lower at baseline in both severely depressed (log_2_ fold difference (LFD)= -0.48; *p*=0.021) and highly anxious (LFD= -0.43; *p*=0.021) participants compared to participants with less severe symptoms. The gut bacteria-derived secondary BAs produced from CDCA such as lithocholic acid (LCA) and several of its metabolites, and their ratios to primary BAs, were significantly higher in the more anxious participants (LFD’s range=[0.23,1.36]; *p*’s range=[6.85E-6,1.86E-2]). The interaction analysis of HRSD_17_ and HRSA-Total suggested that the BA concentration differences were more strongly correlated to the symptoms of anxiety than depression. Significant differences in baseline CDCA (LFD= -0.87, *p*=0.0009), isoLCA (LFD= -1.08, *p*=0.016) and several BA ratios (LFD’s range [0.46, 1.66], *p*’s range [0.0003, 0.049]) differentiated treatment failures from remitters.

**Conclusion:** In patients with MDD, BA profiles representing changes in gut microbiome compositions are associated with higher levels of anxiety and increased probability of first-line treatment failure. If confirmed, these findings suggest the possibility of developing gut microbiome-directed therapies for MDD characterized by gut dysbiosis.

## 1. INTRODUCTION

Abnormalities in the gut microbiome and gut-brain axis have emerged as potentially important contributors to the pathophysiology of neuropsychiatric diseases. Several microbe-derived metabolites (e.g., neurotransmitters, short-chain fatty acids, indoles, bile acids [BAs], choline metabolites, lactate, and vitamins) play a significant role in the context of emotional and behavioral changes (Caspani, Kennedy et al. 2019). Both direct and indirect mechanisms have been proposed through which gut microbial metabolites can affect central nervous system (CNS) functions (Yarandi, Peterson et al. 2016, Tognini 2017, Tremlett, Bauer et al. 2017, Caspani, Kennedy et al. 2019). These include activation of afferent vagal nerve fibers, stimulation of the mucosal immune system or circulatory immune cells after translocation from the gut into the circulation, and absorption into the bloodstream followed by uptake and biochemical interaction with a number of distal organs. In the brain, these metabolites may activate receptors on neurons or glia, modulate neuronal excitability, and change gene expression patterns via epigenetic mechanisms (Caspani, Kennedy et al. 2019).

A growing body of evidence indicates the various mechanisms related to bidirectional communication between the gut microbiota and the host’s CNS with anxiety and depression (Dinan and Cryan 2015, Dinan and Cryan 2017, Rieder, Wisniewski et al. 2017, Simpson, Diaz-Arteche et al. 2021). Certain gut bacteria regulate the production of neurotransmitters and their precursors, such as serotonin, gamma-aminobutyric acid and tryptophan, and they also regulate proteins such as brain-derived neurotrophic factor, a key molecule involved in neuroplastic changes in learning and memory (Bercik, Verdu et al. 2010, O’Sullivan, Barrett et al. 2011, Agus, Planchais et al. 2018, Miranda, Morici et al. 2019). Metabolites such as short-chain fatty acids (Parada Venegas, De la Fuente et al. 2019) are involved in neuropeptide and gut hormone release, and they modulate immune signaling along the gut-brain axis via cytokine production. Gut bacteria are thought to be involved in the development and functioning of the hypothalamic-pituitary-adrenal axis (Sudo, Chida et al. 2004, de Weerth 2017, Foster, Rinaman et al. 2017). Dysregulation of the hypothalamic-pituitary-adrenal axis has been implicated in anxiety and depressive disorders, being associated with higher cortisol levels, increased intestinal permeability, and a sustained proinflammatory state (Keller, Gomez et al. 2017). Gastrointestinal conditions believed to involve gut-microbial dysbiosis and intestinal permeability, such as irritable bowel syndrome, co-occur at remarkably high rates with psychiatric disorders (Simpson, Schwartz et al. 2020). In addition, several animal studies have supported the possibility of gut sysbiosis having a causative role in depression-like behaviors. For example, mice exposed to antibiotics showed gut dysbiosis, depression-like behavior, and altered neuronal hippocampal firing, with reversal of this phenotype following probiotic treatment (Guida, Turco et al. 2018). Transplantation of gut microbiota from humans with major depressive disorder (MDD) to germ-free or microbiota-deficient rodents resulted in a depression-like phenotype, including anhedonia and anxiety-like behaviors (Kelly, Borre et al. 2016, Zheng, Zeng et al. 2016). Despite the literature supporting the involvement of the microbiota-gut-brain axis in mental health disorders, the underlying mechanisms of bidirectional communication and the metabolite mediators by which the gut bacteria regulate the gut-brain connection are not fully understood. Therefore, characterizing the rich array of compounds produced by gut bacteria and defining their protective and cytotoxic effects on the CNS can effectively define targeted interventions.

A potential mechanism by which the gut microbiome may alter CNS function is its impact on BAs. BAs are the amphipathic end products of cholesterol metabolism and can contribute significantly to hepatic, intestinal, and metabolic disorders (Li and Chiang 2014). **Figure 1** shows how BAs are synthesized from cholesterol in the liver via two major pathways, the classical and the alternative; secondary BAs are metabolized by colonic bacteria through multiple and well-characterized enzymatic pathways (Lefebvre, Cariou et al. 2009). Primary BAs are the direct products of cholesterol metabolites in hepatocytes, such as cholic acid (CA) and chenodeoxycholic acid (CDCA). In response to cholecystokinin after feeding, primary BAs are secreted by the liver into the small intestine to ensure absorption of dietary lipids. Accordingly, 95% of the BAs are actively absorbed in the terminal ileum and redirected into the portal circulation to reenter the liver. A small proportion pass into the colon where bacteria transform them into secondary BAs—lithocholic acid (LCA), deoxycholic acid (DCA), and ursodeoxycholic acid—via deconjugation and 7α-dehydroxylation (Hofmann and Hagey 2008, Bajor, Gillberg et al. 2010). Although DCA and LCA are the most abundant secondary BAs, approximately 50 different secondary BAs have been detected in human feces (Devlin and Fischbach 2015).

**Figure 1.**
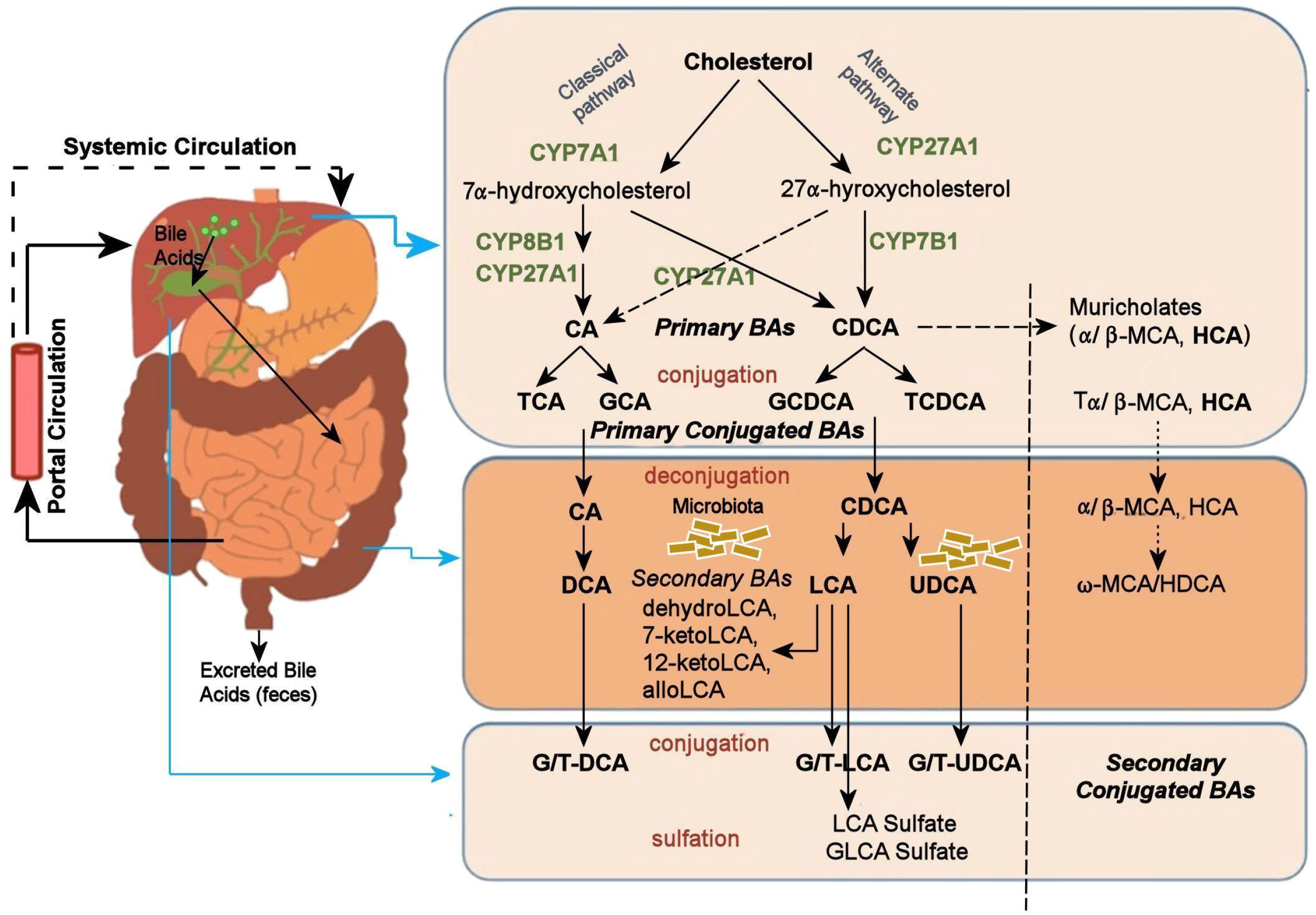
Bile Acid Metabolism Pathway. Bile acids are synthesized from cholesterol in the liver mainly by two pathways. The classical pathway is initiated by the rate-limiting enzyme, CYP7A1 that synthesizes the two primary bile acids in humans, CA and CDCA. CYP8B1 is required for CA synthesis along with the mitochondrial CYP27A1 that catalyzes a steroid side-chain oxidation. The alternative pathway is initiated by CYP27A1, followed by CYP7B1. After synthesis, the primary bile acids are conjugated to the amino acids taurine or glycine for biliary secretion. In the distal ileum and colon, gut bacteria deconjugates the conjugated bile acids, and bacterial 7α-dehydroxylase removes the 7α-hydroxyl group to convert CA and CDCA to the secondary bile acids DCA and LCA, respectively. The LCA as a high toxic bile acid is mostly excreted by feces. A small amount of LCA, which is recycled back into the liver, is subjected to sulfoconjugation at the 3–hydroxy position of sulfotransferase 2A1 (SULT2A1). Sulfoconjugated BAs are almost never reabsorbed by the most important transport proteins, and they are excreted from the body. Several other bacterial modifications are now known that result in the production of a no of different secondary BAs. The classical pathway is the major pathway for daily synthesis of about 80% −90% of the bile acids in humans, whereas the alternative pathway synthesizes about 10-20%. Most bile acids (~95%) are reabsorbed in the ileum and transported via portal blood to the liver to inhibit bile acid synthesis. A small amount of bile acids (~5%) lost in feces is replenished by de novo synthesi *Abbreviations:* ASBT: Apical Sodium-dependent Bile acid Transporters. BA: Bile Acid. BSEP: Bile Salt Export Pump. CA: Cholic Acid. CDCA: Chenodeoxycholic Acid. DCA: Deoxycholic Acid. FXR: Farnesoid X Receptor. GCA: Glycocholic Acid. GCDCA: Glycochenodeoxycholic Acid. GLCA: Glycolithocholic Acid. HCA: Hydroxycitric Acid. HDCA: Hyodeoxycholic Acid. LCA: Lithocholic Acid. MCA: Monocarboxylic Acid. NTCP: Sodium/Taurocholate Co-transporting Polypeptide. SHP: Small heterodimer partner. TCA: Taurocholic Acid. TCDCA: Taurochenodeoxycholic Acid. UDCA: Ursodeoxycholic Acid.

Although primary BAs like CDCA may be synthesized in the brain, no evidence so far supports the synthesis of secondary BAs in the brain (Baloni, Funk et al. 2020). This suggests that the major source of brain BAs is the systemic circulation, which functions as a direct communication bridge between the gut microbiome and the brain (Monteiro-Cardoso, Corliano et al. 2021), thereby playing a vital role in brain health. Circulating BAs generated in the liver and intestine can reach the brain by crossing the blood-brain barrier, either by simple diffusion or through BA transporters (Monteiro-Cardoso, Corliano et al. 2021). Higashi et al. (Higashi, Watanabe et al. 2017) recently found that levels of CA, CDCA, and DCA detected in the brain positively correlated with their serum levels. The liver-gut-brain axis is critical for the maintenance of metabolic homeostasis, yet much remains to be elucidated about how BAs that are synthesized in the liver and modified in the gut mediate the crosstalk between the peripheral and central nervous system and impact neuropsychiatric disorders like depression and anxiety.

Several lines of evidence implicate secondary BAs as contributors to CNS dysfunction. Hepatic encephalopathy is associated with elevated levels of ammonia and cytotoxic BAs, including several conjugated primary and secondary BAs (Xie, Wang et al. 2018). Post-mortem brain samples and serum concentrations of living Alzheimer’s disease patients (compared to health controls) demonstrated lower levels of the primary bile acid, CA, and higher levels of its bacterially-derived secondary bile acid, DCA and its conjugated forms (MahmoudianDehkordi, Arnold et al. 2019, Nho, Kueider-Paisley et al. 2019, Baloni, Funk et al. 2020). In contrast, ursodeoxycholic acid, the 7β isomer of CDCA, has antiapoptotic, anti-inflammatory, antioxidant, and neuroprotective effects in various models of neurodegenerative diseases (Daruich, Picard et al. 2019) (Ramalho, Nunes et al. 2013) and Huntington’s disease (Rodrigues, Stieers et al. 2000) (Mortiboys, Aasly et al. 2013) (Parry, Rodrigues et al. 2010) (Ackerman and Gerhard 2016). Taken together, these data indicate that BAs affect brain function under both normal and pathological conditions. However, the association of BAs on psychiatric diseases such as MDD has received little study to date.

In this study, we profiled baseline serum samples from 208 patients enrolled in a randomized controlled trial of treatment-naïve outpatients with MDD, measuring 36 primary and secondary BAs to address the following questions:

1. Is there a relationship between BA profiles and depressive and anxiety symptom severity?
2. Does symptom severity correlate with differential metabolism of BAs through the classical and alternate pathways?
3. Do baseline BA profiles distinguish MDD patients who achieved remission from those who failed to benefit after 12 weeks of treatment?

## 2. MATERIALS AND METHODS

### 2.1 Study Design and Participants

This study examined serum samples from the Predictors of Remission in Depression to Individual and Combined Treatments (PReDICT) study. The design and clinical outcomes of PReDICT have been detailed previously (Dunlop, Binder et al. 2012, Dunlop, Kelley et al. 2017, Dunlop, LoParo et al. 2019). PReDICT aimed to identify predictors and moderators of response to 12 weeks of randomly assigned treatment with duloxetine (30-60 mg/day), escitalopram (10-20 mg/day) or cognitive behavior therapy (16 one-hour individual sessions). Eligible participants were adults aged 18-65 with nonpsychotic MDD who had never previously been treated for depression. Severity of depression at the randomization visit was assessed with the 17-item Hamilton Depression Rating Scale (HRSD_17_) (Hamilton 1960). Eligibility required an HRSD_17_ score ≥18 at the screening visit and ≥15 at the randomization visit, indicative of moderate-to-severe depression. Patients were excluded if they had a history of bipolar disorder, neurocognitive disorder, or anorexia nervosa, or had an active significant suicide risk, current illicit drug use (assessed by history and with urine drug screen) or a history of substance abuse in the three months prior to randomization, pregnancy, lactation, or any uncontrolled general medical condition.

### 2.2 Metabolomic Profiling and Ratios and Summations

At the randomization visit, antecubital phlebotomy was performed without regard for time of day or fasting status to obtain the serum samples used in the current analysis. Blood samples were allowed to clot for 20 minutes, then centrifuged at 4□C for 10 minutes. The serum was pipetted into Eppendorf tubes and immediately frozen at −80⩑C until ready for metabolomic analysis. Using targeted metabolomics protocols and profiling protocols established in previous studies (Qiu, Cai et al. 2009, Xie, Wang et al. 2015, Zhao, Ni et al. 2017), BAs were quantified by ultra-performance liquid chromatography triple quadrupole mass spectrometry (Waters XEVO TQ-S, Milford, USA). Measures of primary and secondary BAs, including their conjugated and unconjugated forms, can be found in **Supplementary Table 1.**

We examined individual BAs as well as a number of BA summations and ratios that have been previously implicated in several pathophysiological conditions (O’Byrne, Hunt et al. 2003, Shonsey, Sfakianos et al. 2005, Sonne, Hansen et al. 2014, Wahlstrom, Sayin et al. 2016, Chiang 2017, Martinot, Sedes et al. 2017, Vaz and Ferdinandusse 2017, Marksteiner, Blasko et al. 2018, MahmoudianDehkordi, Arnold et al. 2019). See **Supplementary Table 2** for these ratios and their associated diseases or metabolic conditions.

### 2.3 Depression and Anxiety Symptoms

Depression severity was assessed using the clinician-administered HRSD_17_. Participants with HRSD_17_ <20 were labeled as non-severely depressed and those with HRSD_17_ ≥20 as severely depressed (Weitz, Hollon et al. 2015). Anxiety symptom severity was assessed using the clinician-rated 14-item Hamilton Anxiety Rating Scale (HRSA-Total) (Hamilton 1959), comprising two subscales: “psychic anxiety” (items 1–6 and 14) (HRSA-PSY), and “somatic anxiety” (items 7–13) (HRSA-SOM) (Dunlop, Still et al. 2020). Psychic anxiety (HRSA-PSY) consists of the symptoms of anxious mood, tension, fears, depressed mood, insomnia, impaired concentration, and restlessness. Somatic anxiety (HRSA-SOM) consists of physical symptoms associated with the muscular, sensory, cardiovascular, respiratory, gastrointestinal, genitourinary, and autonomic systems. Participants were divided into those with high (HRSA-Total ≥15) and low (HRSA-Total <15) levels of anxiety (Matza, Morlock et al. 2010). The HRSD_17_, and HRSA-Total ratings were re-administered after the completion of treatment at week 12. Consistent with other studies evaluating the biological effects of treatments, we compared the participants who achieved remission (remitters) (defined as completing 12 weeks of treatment and reaching HRSD_17_ ≤7) versus those who completed 12 weeks of treatment but whose week 12 HRSD_17_ score was <30% lower than their baseline score (treatment failure) (Dunlop, Rajendra et al. 2017).

### 2.4 Statistical Analysis

Differences in demographic variables and depression scores across the response groups were evaluated using ANOVA and the Pearson Chi-squared test (for categorical variables). All analyses were performed in a metabolite-wise manner in two ways. 1) Difference in metabolite concentrations in severe vs. non-severe depression, high vs. low anxiety levels, and remission vs. treatment failure were analyzed using the nonparametric, two-sample Wilcoxon signed-rank test. 2) Partial correlations between metabolite levels and the continuous variables HRSD_17_, HRSA-Total, HRSA-SOM, and HRSA-PSY were conducted using partial Spearman rank correlation and adjusted for age, sex, and body mass index. A p-value <0.05 was considered significant. Given the exploratory nature of this initial investigation, no correction for multiple comparisons was made.

We conducted separate partial least squares regression and partial least squares discriminant analysis to examine the contribution of baseline BA levels to baseline HRSD_17_, HRSA-Total, and treatment outcome. In all models, we accounted for age, sex and body mass index, and used 5-fold cross-validation with 100 repeats. In partial least squares regression models, baseline BA profiles of all participants were considered as predictor variables, and the HRSD_17_ and HRSA-Total as continuous dependent variables. Using a partial least squares discriminant analysis model, we examined whether the baseline BA profiles could discriminate participants at the two extremes of the treatment response spectrum, the remitters and those with treatment failure. Significant predictors were identified based on their variable importance on projection scores. Variables with a variable importance on projection score value >1 were considered important for the models.

## 3. RESULTS

### 3.1 Participant Characteristics (Demographic and Clinical)

**Table 1** summarizes the demographic and clinical features of the 208 participants in the PReDICT Study. Of these, 38.94% of participants were male, and mean (standard error of mean) age, HRSD_17_, and HRSA-Total were 38.99(0.81), 19.89(0.26), and 16.40(0.37), respectively. Baseline total HRSD_17_ scores were highly correlated with HRSA-Total scores (Spearman rank correlation *rho=* 0.64) and HRSA-PSY scores (*rho*=0.58), but less strongly correlated with HRSA-SOM (*rho*=0.41). The correlation between HRSA-PSY and HRSA-SOM was only *rho*=0.35 (**Supplementary Figure 1**). Results of PLS regression analyses are presented in **Supplementary Methods and Results**.

**Table 1:** Participant Demographic and Clinical Characteristics.

| Characteristic | Population<br>(N=208) | Depression |  | Anxiety |  | Treatment Outcome |  |
| --- | --- | --- | --- | --- | --- | --- | --- |
|  |  | Non-Severe<br>(N=102) | Severe<br>(N=106) | Low (N=91) | High (N=117) | Remission<br>(N=73) | Treatment<br>Failure<br>(N=25) |
| Age (yrs) <sup>a</sup> | 38.99(0.81) | 36.93(1.14) | 40.97(1.13) | 38.77(1.27) | 46(50.55) | 37.40(1.24) | 37.68(2.61) |
| Sex: Male <sup>b</sup> | 81(38.94 %) | 46(45.10%) | 35(33.02%) | 46(50.55%) | 35(29.91%) | 32(43.84%) | 10(40%) |
| Body Mass Index<br>(kg/m<sup>2</sup>) <sup>a</sup> | 28.78(0.42) | 28.59(0.65) | 28.97(0.55) | 28.34(0.69) | 29.13(0.53) | 29.18(0.75) | 27.62(1.14) |
| HRSD<sub>17</sub> <sup>a</sup> | 19.89(0.26) | 16.82(0.14) | 22.84(0.28) | 17.69(0.29) | 21.60(0.34) | 18.56(0.42) | 19.20(0.69) |
| HRSA-Total <sup>a</sup> | 16.40(0.37) | 13.46(0.38) | 19.24(0.49) | 11.77(0.21) | 20.01(0.39) | 14.78(0.55) | 15.80(1.09) |
| HRSA-SOM <sup>a</sup> | 4.04(0.23) | 2.77(0.26) | 5.25(0.34) | 1.64(0.16) | 5.91(0.29) | 3.21(0.34) | 3.88(0.78) |
| HRSA-PSY <sup>a</sup> | 10.84(0.19) | 9.50(0.21) | 12.12(0.25) | 8.98(0.19) | 12.28(0.22) | 10.04(0.26) | 10.48(0.52) |

<sup>a</sup> Mean and standard error of the mean for each group
<sup>b</sup> Number and percent of males for each group
*Abbreviations:* HRSA-SOM: Somatic anxiety subscore of the Hamilton Anxiety Rating Scale. HRSA-PSY: Psychic anxiety subscore of the Hamilton Anxiety Rating Scale. HRSA-Total: 14-item Hamilton Anxiety Rating Scale. HRSD<sub>17</sub>: 17-item Hamilton Depression Rating Scale.

### 3.2 BA Profiles and Disease Severity

#### 3.2.1. BA Profiles Related to Depressive Symptom Severity

The concentrations of the conjugated and unconjugated versions of the primary and secondary BAs are reported in **Supplementary Table 1.**

##### 3.2.1.1. Primary BAs

As depicted in **Figure 2**, the primary bile acid CDCA, which is produced predominantly from the alternate pathway, was negatively correlated with the baseline total HRSD_17_ score after adjusting for age, sex, and body mass index (partial correlation *rho*= −0.16, *p*=0.021). Dichotomous analysis showed a significantly lower CDCA in the more compared to the less severely depressed participants (LFD= −0.48, *p*=0.02). No significant correlation or difference was noted for CA, the primary BA produced through the classical pathway (*rho*= −0.01, *p*=0.88; *p*_Wilcoxon_=0.41).

**Figure 2:**
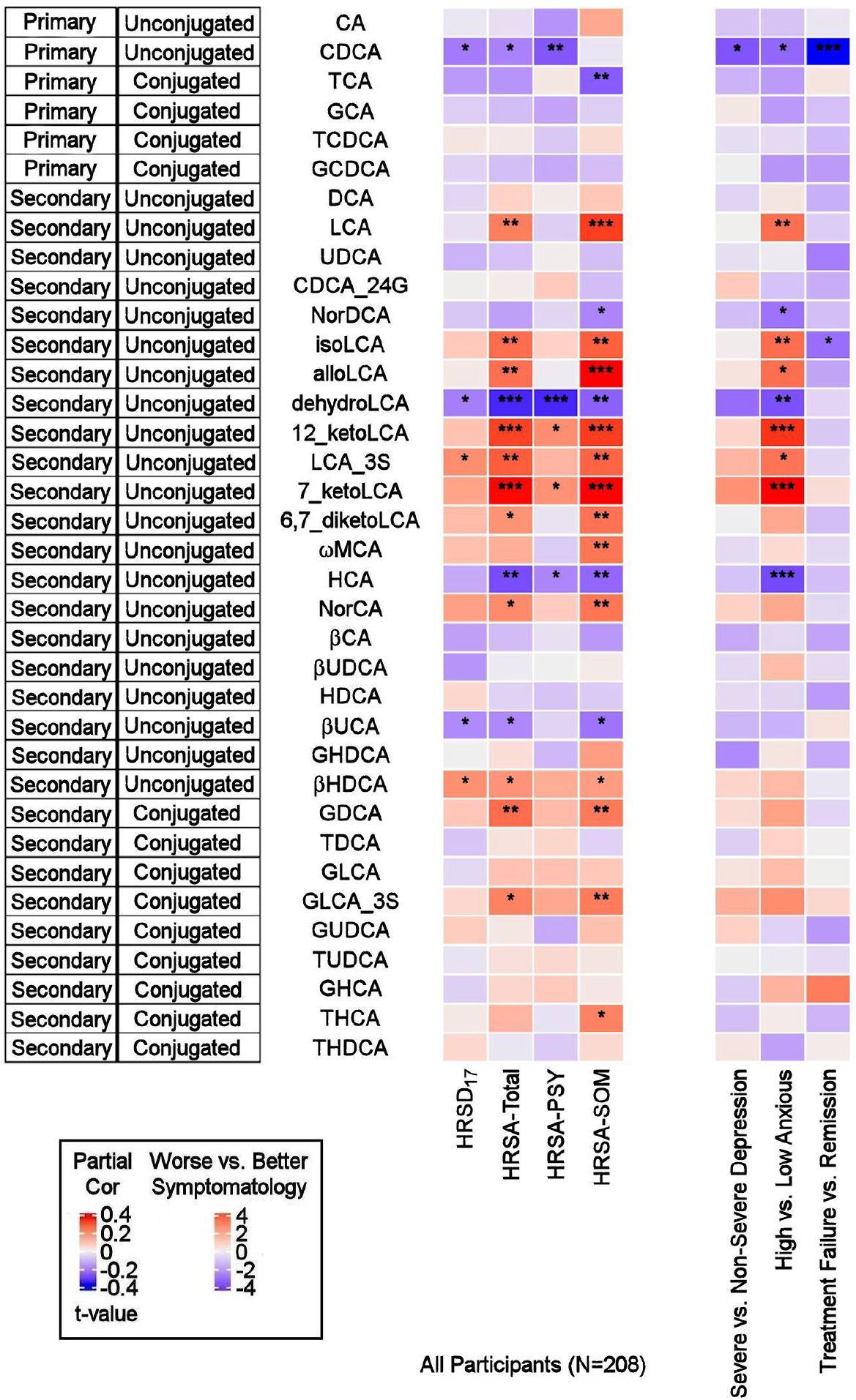
Correlations between Baseline BAs and Depression and Anxiety Scores, and Differences in Baseline BA Profiles between Several Participant Groups. On the left: Heat map of partial Spearman rank correlations between baseline BAs and scores on the HRSD_17_ and Hamilton Anxiety Rating Scale and subscales, after accounting for age, sex, and body mass index. On the right: Heat map of differences in baseline BA profiles in severe vs. non-severe depressed, high vs. low anxiety and treatment-failure vs. remitter groups. T-values were used for visualization purposes and the Wilcoxon Ranked Sum Test were used to test the significance of differences. *Abbreviations:* BA: Bile Acid. CA: Cholic Acid. CDCA: Chenodeoxycholic Acid. DCA: Deoxycholic Acid. GCA: Glycocholic Acid. GCDCA: Glycochenodeoxycholic Acid. GDCA: Glycodeoxycholic Acid. GHCA: Glycohyocholic Acid. GHDCA: Glycohyodeoxycholic Acid. GLCA: Glycolithocholic Acid. GUDCA: Glycoursodeoxycholic Acid. HCA: Hydroxycitric Acid. HDCA: Hyodeoxycholic Acid. HRSA-PSY: Psychic anxiety subscore of the Hamilton Anxiety Rating Scale. HRSA-SOM: Somatic anxiety subscore of the Hamilton Anxiety Rating Scale. HRSA-Total: 14-item Hamilton Anxiety Rating Scale. HRSD_17_: 17-item Hamilton Depression Rating Scale. LCA: Lithocholic Acid. MCA: Monocarboxylic Acid. TCA: Taurocholic Acid. TCDCA: Taurochenodeoxycholic Acid. TDCA: Taurodeoxycholic Acid. THCA: Tetrahydrocannabinolic Acid. THDCA: Taurohyodeoxycholic Acid. TUDCA: Tauroursodeoxycholic Acid. UCA: Ursocholic Acid. UDCA: Ursodeoxycholic Acid. _3S: 3 Sulfate. *: uncorrected p-value<0.05. **: uncorrected p-value< 0.01. ***: uncorrected p-value< 0.001.

##### 3.2.1.2. Secondary BAs

The secondary bacterially-produced BAs, lithocholic acid 3 sulfate (LCA_3S) and isohyodeoxycholic acid (βHDCA) were positively correlated with HRSD_17_ (*rho*=0.158, *p*=0.022 and rho=0.156, p=0.025, respectively) while dehydro-LCA was negatively correlated (*rho*= −0.154, *p*=0.027). Similar trends were noted in non-severe vs. severe depressed groups for the aforementioned analytes, but the differences did not reach the significance level.

#### 3.2.2. BA Profiles Related to Anxiety Symptom Severity

##### 3.2.2.1. Primary BAs

CDCA was negatively correlated with HRSA-Total (*rho*= −0.149, *p*=0.032) and HRSA-PSY (*rho*= −0.207, *p*=0.0028), but not HRSA-SOM (*rho*= −0.015, *p*=0.82). CDCA was significantly lower in the highly anxious participants (*p*=0.021). No significant correlation was noted for the other primary bile acid, CA (classical pathway). However, norcholic acid, which is a non-conjugated C23 homologue of the primary bile acid, CA, exhibited positive correlations with HRSA-Total (*rho*= 0.163, *p*=0.019), and HRSA-SOM (*rho*= 0.195, *p*=0.015).

##### 3.2.2.2. Secondary BAs

The bacterially derived 7β-hydroxy epimer of CA, β-ursocholic acid and the CDCA-derived hyocholic acid were inversely correlated with HRSA-Total and HRSA-SOM (*rho*’s range [−0.22 to −0.13], *p*’s range [0.001 to 0.046]). LCA, produced by 7-alpha-dehydroxylation of CDCA, and several of its derivatives including 7-keto-LCA, isoLCA, alloLCA, and 12-ketoLCA, were strongly positively correlated with HRSA-Total and HRSA-SOM (*rho*’s range [0.18-0.34], *p*’s range [4.46E-07 to 8.85E-03]). These BAs were also significantly elevated or trended to be elevated in highly anxious compared to less anxious participants (*p*’s between 0.0002-0.01). In contrast to LCA and many of its derivatives that correlated positively with anxiety severity, dehydroLCA (a known anti-inflammatory BA) was negatively correlated with HRSA-Total (*rho*= −0.266, *p*=0.0001), HRSA-SOM (*rho*= −0.195, *p*=0.004) and HRSA-PSY (*rho*= −0.266, *p*=0.0001). In addition, two secondary glycine conjugated BAs were positively correlated with HRSA-Total and HRSA-SOM scores: glycodeoxycholic acid (GDCA) (HRSA-Total: *rho*=0.20, *p*=0.002; HRSA-SOM: *rho*=0.18, *p*=0.006) and glycolithocholic acid 3 sulfate (GLCA_3S) (HRSA-Total: *rho*=0.17, *p*=0.011; HRSA-SOM: *rho*=0.187, *p*=0.007).

Overall, greater baseline anxiety was associated with *lower* concentrations of the primary BAs (primarily CDCA) and their conjugated forms, and *higher* levels or concentrations of secondary BAs, derived from CDCA, such as the hepatotoxic LCA and many of its metabolites. The correlations between the secondary BAs and HRSA-Total score were driven primarily by somatic anxiety symptoms.

To investigate whether the differences observed in the BAs reported above were driven by anxiety or depression, we further tested the interaction effect of severity of anxiety and depression on the BAs. As shown in **Figure 3**, several gut-microbe-produced BAs and ratios of secondary to primary BAs (e.g., LCA, 7-ketoLCA, 12-ketoLCA, LCA/CDCA, 7-ketoLCA/CDCA, alloLCA/CDCA, 12-ketoLCA/CDCA) significantly differed between low versus highly anxious MDD participants irrespective of depression severity. For example, LCA levels were significantly higher in both non-severe depression-high anxiety and high depression-high anxiety participants compared to the non-severe depression-low anxiety and severe depression-low anxiety groups respectively (*p*=0.012 and p=0.016, respectively). This was also observed with the other CDCA derived BAs or the ratios (**Figure 3)**. These data suggest that the differences in these BA profiles are significantly associated with anxiety but not depressive symptom severity.

**Figure 3:**
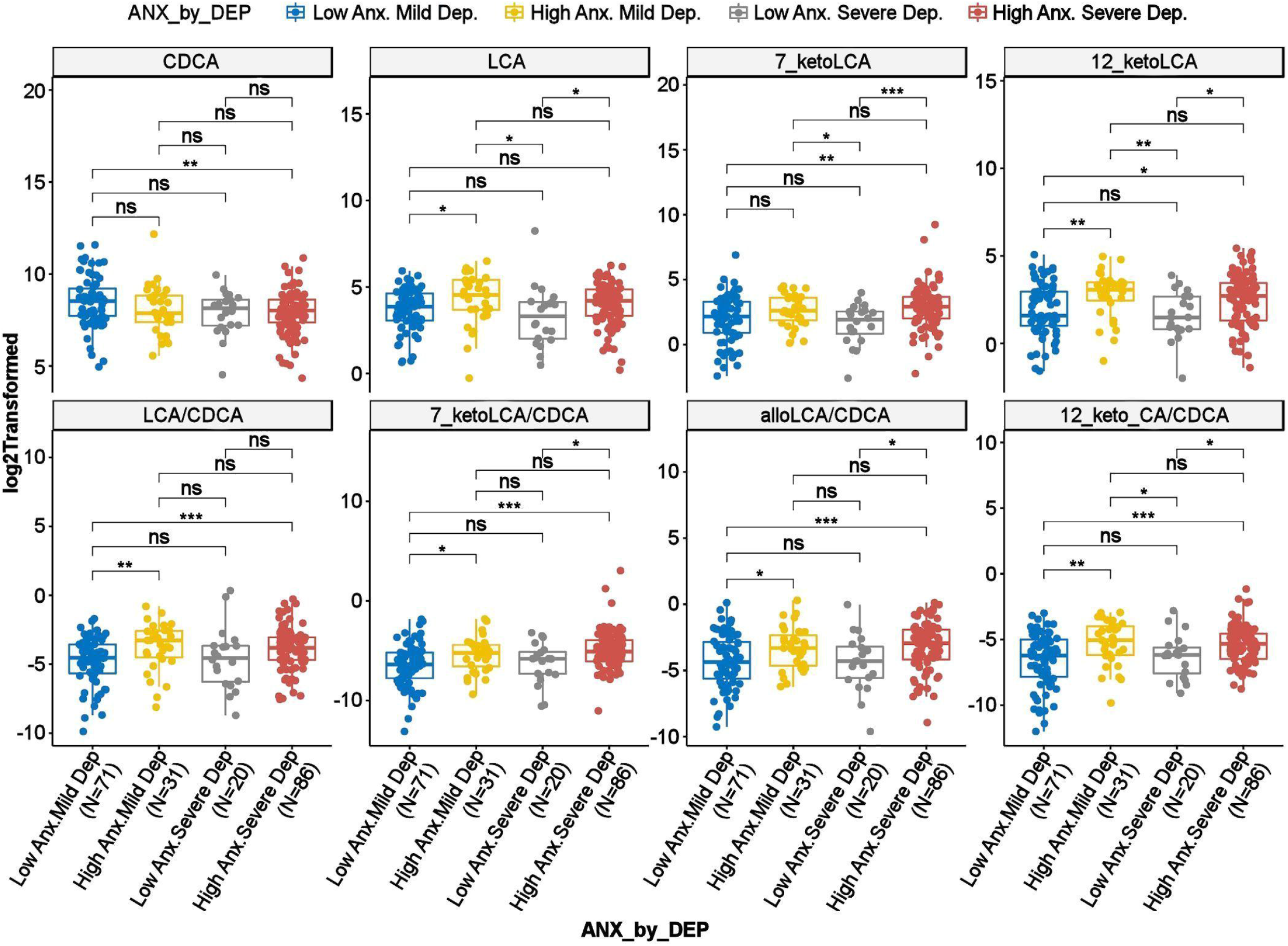
Scatter Plots of HRSD□ Scores by HRSA-Total Interaction for Selected Bile Acids and Ratios. *Abbreviations:* Anx: Anxiety. CA: Cholic Acid. CDCA: Chenodeoxycholic Acid. Dep: Depression. HRSA-Total: 14-item Hamilton Anxiety Rating Scale. HRSD_17_: 17-item Hamilton Depression Rating Scale. LCA: Lithocholic Acid. *: uncorrected p-value<0.05. **: uncorrected p-value< 0.01. ***: uncorrected p-value< 0.001; ns: not significant

### 3.3 Altered Metabolism of BAs through Classical and Alternate Pathways in MDD participants

To investigate potential shifts in BA synthesis pathways or possible alterations in enzymatic activities, we further examined all possible pairwise BA ratios and selected composite summations and ratios that can inform about changes in classical and alternate pathways of BA metabolism. A list of the BA summations and ratios and their implicated pathophysiology are shown in **Supplementary Table 2**. Partial correlation analysis of depression severity score with composite summations and ratios did not yield strong correlation (**Figure 4A**). However, a few ratios showed significant differences between participants with non-severe versus severe symptoms of anxiety. A higher value of the ratio of “primary BAs to total BAs,” which represents a fraction of primary BAs relative to the BA pool, was correlated to less severe anxiety. Concomitantly, lower values of the “secondary to primary BAs” ratio, which represents a fraction of secondary BAs relative to the BA pool, as well as “Secondary BA Synthesis”, which is the ratio of cytotoxic secondary BAs to primary BAs, were correlated with less severe anxiety symptomology (HRSA-Total). Both HRSA-PSY and HRSA-SOM were similarly affected (absolute *rho*’s range [0.19 to 0.25],*p’s* range [2.14E-4 to 5.11E-3]). Additionally, “sum of unconjugated primary BAs”, a higher level of which may indicate less BA conjugation and less solubility, was negatively correlated with HRSA-PSY (*rho*=-0.22, *p*=9.61E-4).

**Figure 4:**
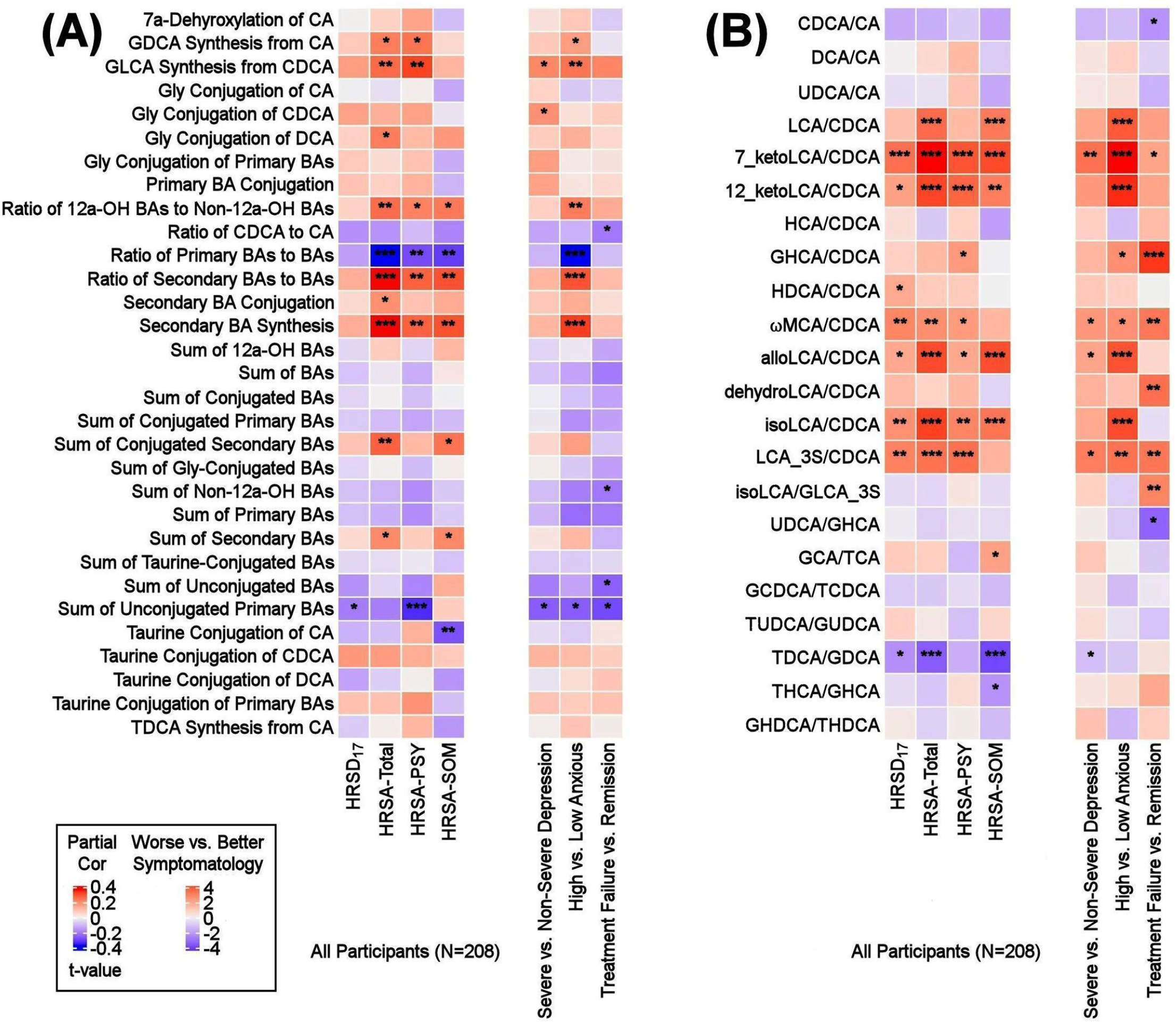
Ratios of BAs Reflective of Liver and Gut Microbiome Enzymatic Activities in Depressed Patients. Three types of ratios (pairwise or composite) were calculated to inform about possible enzymatic activity changes in depressed participants. These ratios reflect one of the following: (1) Shift in BA metabolism from primary to alternative pathway. (2) Changes in gut microbiome correlated with production of secondary BAs. (3) Changes in glycine and taurine conjugation of BAs. (A) Composite Ratios and summations. (B) Selected Pairwise Ratios. For each figure, the left panel presents a heat map of partial Spearman rank correlations between BA ratios/summations and scores on the HRSD_17_ and Hamilton Anxiety scale and subscales, after accounting for age, sex, and body mass index, and the right panel presents a heat map of differences in ratios/summations in severe vs. non-severe depressed, high vs. low anxious and treatment-failure vs. remitter groups. *Abbreviations:* BA: Bile Acid. CA: Cholic Acid. CDCA: Chenodeoxycholic Acid. DCA: Deoxycholic Acid. GCA: Glycocholic Acid. GCDCA: Glycochenodeoxycholic Acid. GDCA: Glycodeoxycholic Acid. GHCA: Glycohyocholic Acid. GHDCA: Glycohyodeoxycholic Acid. GLCA: Glycolithocholic Acid. GUDCA: Glycoursodeoxycholic Acid. HCA: Hydroxycitric Acid. HDCA: Hyodeoxycholic Acid. HRSA-PSY: Psychic anxiety subscore of the Hamilton Anxiety Rating Scale. HRSA-SOM: Somatic anxiety subscore of the Hamilton Anxiety Rating Scale. HRSA-Total: 14-item Hamilton Anxiety Rating Scale. HRSD_17_: 17-item Hamilton Depression Rating Scale (HRSD_17_). LCA: Lithocholic Acid. MCA: Monocarboxylic Acid. TCA: Taurocholic Acid. TCDCA: Taurochenodeoxycholic Acid. TDCA: Taurodeoxycholic Acid. THCA: Tetrahydrocannabinolic Acid. THDCA: Taurohyodeoxycholic Acid. TUDCA: Tauroursodeoxycholic Acid. UDCA: Ursodeoxycholic Acid. *: uncorrected p-value<0.05. **: uncorrected p-value< 0.01. ***: uncorrected p-value< 0.001.

Ratios of CDCA/CA, which is an indicator of a shift in BA synthesis from classical to alternate pathway, as well as conjugated/unconjugated BA ratio for the taurine or glycine conjugations, did not yield significant correlations.

In high anxiety vs. low anxiety participants, the most significant differences in pairwise ratios were observed in the ratios of secondary to the (precursor) primary CDCA such as LCA/CDCA (*p*= 0.0001), 7-ketoLCA/CDCA (*p*=6.85e-06), 12-ketoLCA/CDCA (*p*=4.87e-05), alloLCA/CDCA (*p*=0.0001), isoLCA/CDCA (*p*=3.57e-05), LCA-3S/CDCA (*p*=0.002), glycohyocholic acid (GHCA)/CDCA(*p*=0.041), omega monocarboxylic acid (ωMCA)/CDCA (*p*= 0.021), all of which were significantly higher in participants with more severe symptoms, particularly HRSA-PSY. This suggests an increased utilization of CDCA for the synthesis of bacterially-derived secondary BA in these participants (**Figure 4B**).

Partial correlation analysis of BA ratios and anxiety scores also showed that the gut-bacteria-produced secondary BAs to their precursor primary BA ratios such as LCA/CDCA, 7-ketoLCA/CDCA, 12-ketoLCA/CDCA, alloLCA/CDCA, isoLCA/CDCA, LCA-3S/CDCA were significantly positively correlated with anxiety symptoms (*rho*’s range [0.14 to 0.35], *p*’s range [2.32e-07 to 4.16e-02]). The ratio of the taurine to glycine conjugated deoxycholic acid, TDCA/GDCA, was significantly negatively correlated to HRSA-SOM (*rho*= −0.27; *p*=7.22e-05). Overall, our ratio data indicated a significant trend towards higher levels of secondary BAs compared to their primary precursors that correlated with more anxiety severity in these MDD participants, which suggests gut microbiome dysbiosis in more anxious patients.

### 3.4. Do Baseline BA Concentrations Distinguish Participants who Reached Symptom Remission from those Who Experienced Treatment Failure from 12 Weeks of Treatment?

We further examined whether any of the metabolites that were associated with depression and/or anxiety symptom severity at baseline were different in participants who responded to treatment (remitters; N=73) versus those who did not respond to treatment (treatment failures; N=25) after 12 weeks of treatment/therapy. The metabolites which showed significantly higher baseline levels (*p*<0.05) in remitters compared to the treatment failures were the primary bile acid, CDCA (*p*=0.0009), its bacterial derivative isoLCA (*p*=0.0162) (**Figures 2 and 5**) and the ratio of the two primary bile acids CDCA/CA (*p*=0.0495) (**Figures 4B** and **5**). Several secondary BA to CDCA ratios such as 7-ketoLCA/CDCA, GHCA/CDCA, ωMCA/CDCA, dehydroLCA/CDCA, LCA-3S/CDCA and the secondary to secondary ratio, GLCA-3S/isoLCA (**Figures 4B and 5)** were significantly lower at baseline in the remitters compared to the treatment failures (*p*’s range [0.00032-0.0495]). A summary model of secondary BA synthesis from CDCA and their alterations in these participants is presented in **Figure 6**.

**Figure 5:**
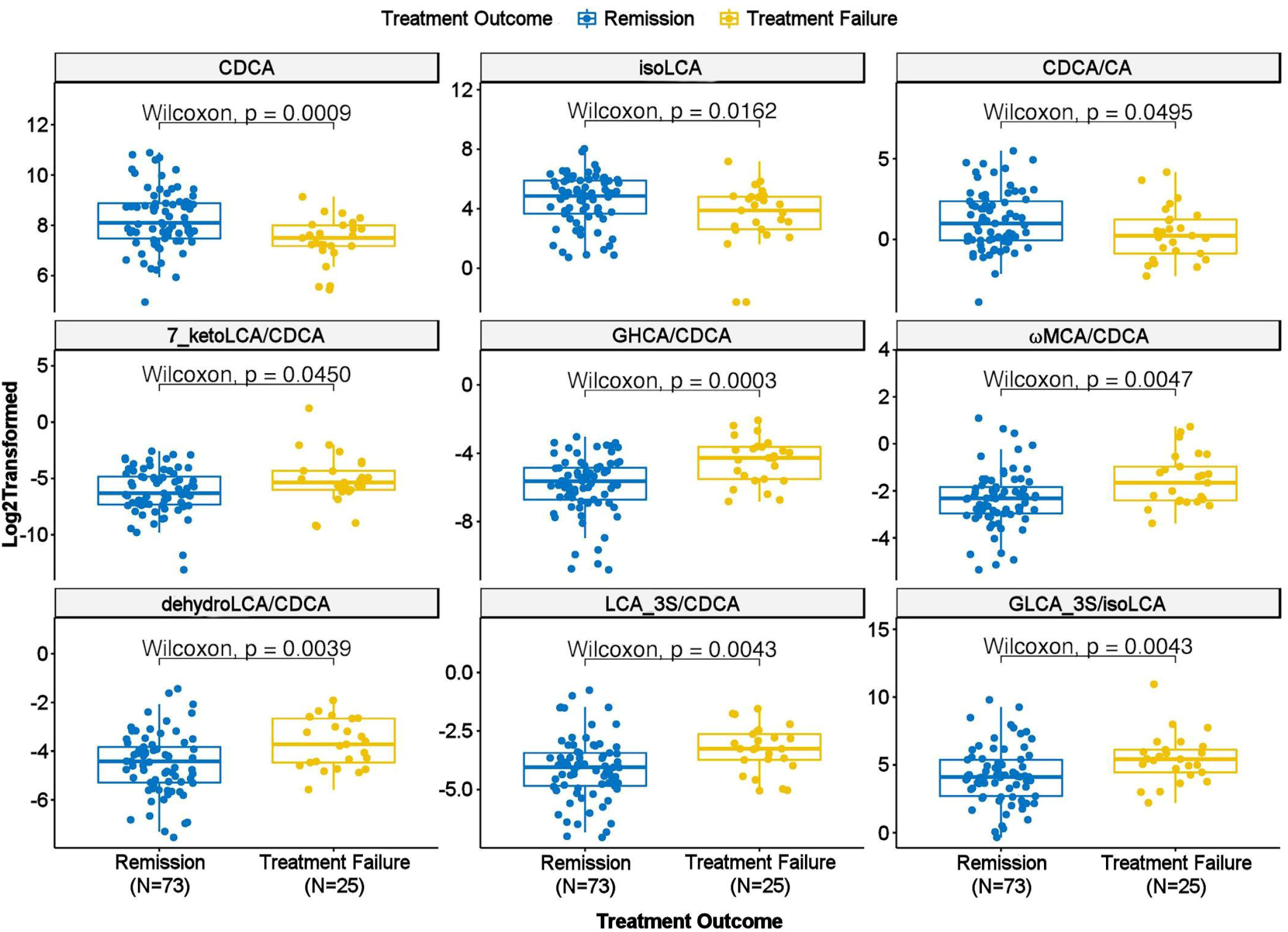
Scatter Plot of Baseline Concentration of Selected Bile Acids and Bile Acid Ratios in Treatment Failure versus Remission Groups. *Abbreviations:* CA: Cholic Acid. CDCA: Chenodeoxycholic Acid. GHCA: Glycohyocholic Acid. GLCA: Glycolithocholic Acid. LCA: Lithocholic Acid. MCA: Monocarboxylic Acid.

**Figure 6:**
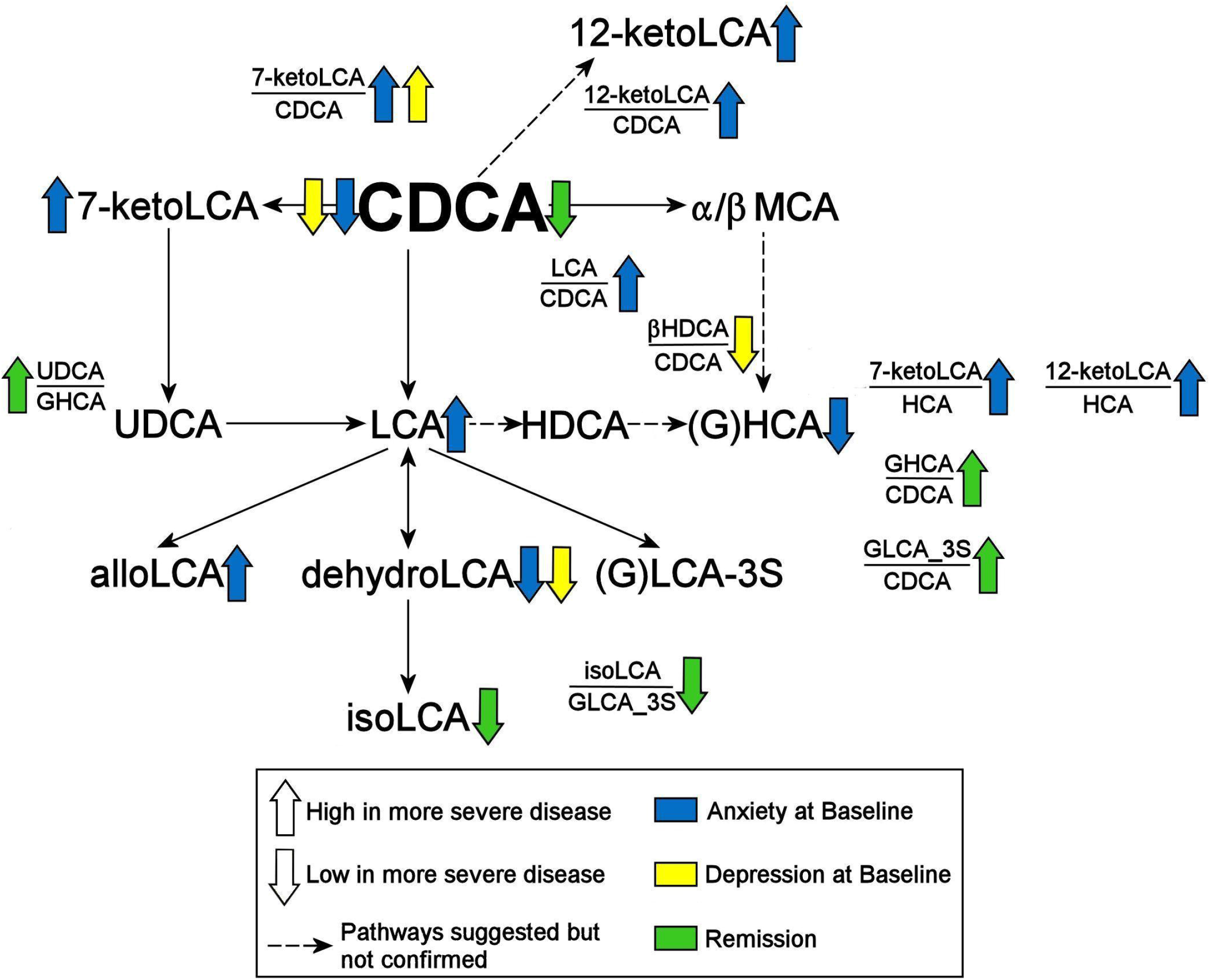
Summary of Findings. *Abbreviations:* CDCA: Chenodeoxycholic Acid. GHCA: Glycohyocholic Acid. GLCA: Glycolithocholic Acid. HCA: Hydroxycitric Acid. HDCA: Hyodeoxycholic Acid. LCA: Lithocholic Acid. MCA: Monocarboxylic Acid. UDCA: Ursodeoxycholic Acid. _3S: 3 Sulfate.

## 4. DISCUSSION

Mounting evidence indicates that gut dysbiosis and the bidirectional communication between brain and gut microflora play an important role in the development of neuropsychiatric diseases. Using targeted metabolomics in participants with MDD, we determined that increased levels of cytotoxic secondary BAs, bacterially-derived from the primary bile acid CDCA, correlated with anxiety symptom severity. The classical pathway that, predominantly, produces the primary bile acid CA seemed to be less impacted. Additionally, participants who did not benefit from treatment were found to have higher baseline levels of the cytotoxic secondary BAs derived from CDCA. Our findings suggest that alternate therapies might be needed that target the gut microbiome for patients who have gut dysbiosis.

We first addressed whether BA concentrations impacted depression and anxiety symptom severity (**Figure 2** and **6**). Overall, BA concentrations appeared to have a stronger impact on anxiety than on depression. Several secondary BA concentrations, and the ratios of secondary to primary BAs, were significantly different between low versus high-anxious MDD participants irrespective of depression severity (**Figure 3**). These secondary BAs included LCA and its derivatives, 7-keto-LCA, isoLCA, alloLCA and 12-ketoLCA. The 7α-dehydroxylation reaction that results in the formation of the secondary BAs has been described as the most quantitatively important process performed by colonic bacteria belonging to the genus *Clostridium,* an enzymatic reaction that is impacted in many neurological diseases (Kiriyama and Nochi 2019). LCA is produced by 7α-dehydroxylation of CDCA and is known to be cytotoxic in rodents as well as several human cell types.

Our second question addressed whether there were any associations of symptoms with the classical and alternate pathways of BA synthesis. In Alzheimer’s disease, we had observed a significant shift in BA synthesis from classical to the alternative pathways in the Alzheimer’s participants compared to healthy controls (MahmoudianDehkordi, Arnold et al. 2019, Nho, Kueider-Paisley et al. 2019, Baloni, Funk et al. 2020). In these MDD participants, we observed that the alternate pathway that favors CDCA synthesis was significantly impacted in the highly-anxious participants. However, no shift from classical to alternate pathway could be observed in these participants since the ratio of CA/CDCA, indicative of such a shift, was not significantly associated with symptom severity (**Figure 4**). Lower CDCA levels and higher secondary metabolites derived from CDCA (and mostly higher ratios of these secondary BAs to CDCA) characterized the participants with higher symptom severity, which may indicate greater utilization of CDCA by the gut bacteria. We also found no significant impact of glycine and taurine conjugation of BA on symptom severity (**Figure 4)**. Interestingly, dehydrolithocholic acid, a major metabolite of LCA, was strongly negatively correlated to anxiety levels in the MDD participants. It is an agonist of the nuclear receptors GPCR1, the farnesoid X receptor (FXR) and the pregnane X receptor, and has recently been shown to regulate adaptive immunity by inhibiting the differentiation of TH17 cells that are known to contribute autoimmunity and inflammation (Hang, Paik et al. 2019). Our final question examined whether any relationship exists between baseline metabolite levels and response to treatment. Remitters showed higher levels of CDCA and one of its gut microbial metabolites (isoLCA) compared to participants for whom treatment failed.

The enzymatic processes involved in altered BA metabolism in CNS diseases may be informed by the association of BAs with inborn errors of metabolism (IEM), in which reduced intestinal BA concentrations result in serious morbidity or mortality. To date, investigators have identified nine recognized IEMs of BAs that lead to enzyme deficiencies and impaired BA synthesis (Heubi, Setchell et al. 2007, Sundaram, Bove et al. 2008). These diseases are characterized by a failure to produce primary BAs and an accumulation of unusual BAs and BA intermediaries. Administration of BAs for replacement therapy often improves the symptoms of IEM, such as cerebrotendinous xanthomatosis, with CDCA the predominant choice for treating both neurological and non-neurological symptoms (Nie, Chen et al. 2014). We have recently reported on a common link between IEM and depression through acylcarnitines and beta oxidation of fatty acids, in which medium-chain acyl-coenzyme A dehydrogenase, an enzyme involved in the production of medium chain acylcarnitines, was shown to be causally linked to depression and also to IEM. These emerging data linking metabolomic disturbances in CNS disorders and IEM provide novel insights into pathobiological processes that contribute to psychiatric disorders (Milaneschi, Arnold et al. 2021).

BAs influence metabolic processes by acting as signaling molecules via the nuclear receptors FXR, the pregnane X receptor, the vitamin D receptor, Takeda G-protein-coupled bile acid receptor, and sphingosine-1-phosphate receptor 2, initiating a variety of signaling cascades relevant to metabolic and hepatic diseases such as obesity, steatosis and steatohepatitis, as well as liver and colon cancer (Lefebvre, Cariou et al. 2009, Wan and Sheng 2018). FXR plays many important roles in the regulation mechanisms of BA synthesis and transport. FXR activation represses the expression of the main enzymes in BA synthesis, CYP7A1 and CYP27A1 (Pauli-Magnus and Meier 2005). In contrast, FXR activation upregulates UGT2B4, which is involved in the conversion of hydrophobic BAs to their less toxic glucuronide derivatives (Barbier, Torra et al. 2003). CDCA is the most potent activator of FXR. Studies in knockout mice suggest the involvement of FXR in modulating brain function. Deletion of FXR altered the levels of several neurotransmitters in the hippocampus and cerebellum, and impaired cognitive functioning and motor coordination (Huang, Wang et al. 2015), which suggests that FXR signaling is required for normal brain function. A recent study using a rat-model (Chen, Zheng et al. 2018) found that over-expression of hippocampal FXR mediated chronic unpredictable stress-induced depression-like behaviors and decreased hippocampal brain-derived neurotrophic factor expression, and that knocking out of hippocampal FXR completely prevented depressive behaviors via brain-derived neurotrophic factor expression.

LCA is the most potent ligand for Takeda G-protein-coupled BA receptor (Kawamata, Fujii et al. 2003), and BA-dependent Takeda G-protein-coupled BA receptor-mediated signaling has been shown to influence the brain by regulating the production of the gut peptide hormone GLP-1 (Monteiro-Cardoso, Corliano et al. 2021), which potentiates glucose-stimulated insulin secretion. LCA is also a potent activator of pregnane X receptor and vitamin D receptor. Thus, largely through their binding and activation of these receptors, BAs regulate their own synthesis, conjugation, transport and detoxification, as well as lipid, glucose, and energy homeostasis (Hylemon, Zhou et al. 2009, Li and Chiang 2015, Ridlon, Harris et al. 2016, Grant and DeMorrow 2020).

The decrease in CDCA with concomitant increase in LCA has particular pathognomonic significance in MDD patients. LCA is formed in humans mainly from the intestinal bacterial 7α-dehydroxylation of CDCA and comprises less than 5% of the total BA pool in humans but is one of the most hydrophobic naturally occurring BAs (Ceryak, Bouscarel et al. 1998).

LCA has been shown to induce double-strand breaks in DNA (Kulkarni, Heidepriem et al. 1980). The mammalian host responds by metabolizing LCA, mainly through sulfation, enabling more efficient excretion and reduced hydrophobicity (Ridlon and Bajaj 2015). BA sulfation is an important detoxification process that converts hydrophobic BAs into excretable metabolites in the liver. Sulfation is catalyzed by a group of enzymes called sulfotransferases (Ridlon and Bajaj 2015). Although, only a small proportion of BAs in bile and serum are sulfated, more than 70% of BAs in urine are sulfated, indicating their efficient elimination in urine (Alnouti 2009). It is estimated that 40–75% of the hydrophobic, hepatotoxic LCA in human bile is present in the sulfated form (Palmer and Bolt 1971). The formation of BA-sulfates increases during cholestatic diseases. Therefore, sulfation may play an important role in maintaining BA homeostasis under pathologic conditions. In our study, we observed elevated levels of the sulfated form of the toxic LCA and GLCA in more severely anxious patients. We have also previously reported increased production of other bacterially-derived sulfates like p-cresol sulfate and indoxyl sulfates (Brydges, Fiehn et al. 2021) in the PReDICT study participants. Together, these findings suggest that alterations in sulfotransferase activities may occur in the liver of some patients.

The microbial conversion of CDCA to 7-keto-LCA, present at higher levels in highly-anxious MDD participants, is known to be reduced in the liver by human 11β-HSDH-1, an enzyme with the primary function of converting cortisone to the active glucocorticoid, cortisol (Odermatt, Da Cunha et al. 2011). Microbial-derived 7-keto-LCA acts as a competitive inhibitor of 11β-HSDH-1, and thus may influence the ratio of cortisone/cortisol.

Among the secondary BAs, dehydrolithocholic acid, a major metabolite of LCA, was interestingly the only metabolite strongly negatively correlated to anxiety levels and depression level as well, in the MDD participants. It is an agonist of the nuclear receptors, GPCR1, FXR, PXR, and has recently been shown to regulate adaptive immunity by inhibiting the differentiation of TH17 cells that are known to cause autoimmunity and inflammation (Hang, Paik et al. 2019).

There are a few limitations to our study. First, we lacked a healthy control group to compare with the participants who had MDD. Second, we did not apply multiple comparison adjustments due to the relatively small sample size and the exploratory nature of this study. Third, these findings will require replication in an independent cohort. Fourth, a number of novel BAs have recently been discovered and were not included in our metabolomic analyses; these compounds should be evaluated in future studies.

It has been suggested (Hibbing, Fuqua et al. 2010, Foster, Schluter et al. 2017) that in the highly evolutionary competitive environment of the human gut microbiome, the persistence of these microbial enzyme activities usually indicates that they increase the organism’s ability to survive. However, dysbiosis in the gut is also possible. Our data suggest that low levels of CDCA might be a result of increased utilization for production of bacterial products in the intestine which, in turn, suggest gut-microbe composition changes or associated enzymatic changes. The underlying pathophysiological significance of BA pool changes remain to be determined, but a reasonable hypothesis emerging from this work is that increases in circulating BAs result from a more hydrophobic BA pool in the colon consequent to gut microbial dysbiosis. These BAs may then produce enhanced toxicity and pathophysiology to cells in the liver, gastrointestinal tract, and the brain.

## Supporting information

Supplemental Results

## AUTHOR CONTRIBUTIONS

SM and SB did analysis of data and helped write the manuscript; CRB did PLS regression analysis. WJ and his team generated biochemical data; RRK, BWD and AJR helped with interpretation of findings and clinical relevance; RKD is PI for project and helped with concept development, study design, data interpretation and connecting biochemical and clinical data, and with the writing of the manuscript.

## ABBREVIATIONS

BA: Bile Acid
CA: Cholic Acid
CDCA: Chenodeoxycholic Acid
CNS: Central Nervous System
DCA: Deoxycholic Acid
FXR: Farnesoid X Receptor
GDCA: Glycodeoxycholic Acid
GHCA: Glycohyocholic Acid
GLCA: Glycolithocholic Acid
HRSA-PSY: Psychic Anxiety subscale of the 14-item Hamilton Anxiety Rating Scale
HRSA-SOM: Somatic Anxiety subscale of the 14-item Hamilton Anxiety Rating Scale
HRSA-Total: 14-item Hamilton Anxiety Rating Scale
HRSD_17_: 17-item Hamilton Depression Rating Scale
IEM: Inborn Errors of Metabolism
LCA: Lithocholic Acid
MCA: Monocarboxylic Acid
MDD: Major Depressive Disorder
PReDICT: Predictors of Remission in Depression to Individual and Combined Treatments study
TDCA: Taurodeoxycholic Acid

## ACKNOWLEDGEMENTS

We acknowledge the editorial services of Mr. Jon Kilner, MS, MA (Pittsburgh) and the assistance of Ms. Lisa Howerton (Duke).

## CONTRIBUTION TO THE FIELD STATEMENT

This study contributes to the field of major depressive disorder by mapping the biochemical changes associated with gut-microbiome dysbiosis in depression. Bile acids which are end products of cholesterol metabolism in the liver are modified in the gut to produce secondary bile acids. Our results provide insights into how the gut-microbiota can impact the severity of anxiety distress and depression as well as treatment response through the altered biosynthesis of these secondary metabolites.

## References

Ackerman, H. D. and G. S. Gerhard (2016). “Bile Acids in Neurodegenerative Disorders.” Front Aging Neurosci 8: 263.

Agus, A., J. Planchais and H. Sokol (2018). “Gut Microbiota Regulation of Tryptophan Metabolism in Health and Disease.” Cell Host Microbe 23(6): 716–724.

Alnouti, Y. (2009). “Bile Acid sulfation: a pathway of bile acid elimination and detoxification.” Toxicol Sci 108(2): 225–246.

Bajor, A., P. G. Gillberg and H. Abrahamsson (2010). “Bile acids: short and long term effects in the intestine.” Scand J Gastroenterol 45(6): 645–664.

Baloni, P., C. C. Funk, J. Yan, J. T. Yurkovich, A. Kueider-Paisley, K. Nho, A. Heinken, W. Jia, S. Mahmoudiandehkordi, G. Louie, A. J. Saykin, M. Arnold, G. Kastenmuller, W. J. Griffiths, I. Thiele, C. Alzheimer’s Disease Metabolomics, R. Kaddurah-Daouk and N. D. Price (2020). “Metabolic Network Analysis Reveals Altered Bile Acid Synthesis and Metabolism in Alzheimer’s Disease.” Cell Rep Med 1(8): 100138.

Barbier, O., I. P. Torra, A. Sirvent, T. Claudel, C. Blanquart, D. Duran-Sandoval, F. Kuipers, V. Kosykh, J. C. Fruchart and B. Staels (2003). “FXR induces the UGT2B4 enzyme in hepatocytes: a potential mechanism of negative feedback control of FXR activity.” Gastroenterology 124(7): 1926–1940.

Bercik, P., E. F. Verdu, J. A. Foster, J. Macri, M. Potter, X. Huang, P. Malinowski, W. Jackson, P. Blennerhassett, K. A. Neufeld, J. Lu, W. I. Khan, I. Corthesy-Theulaz, C. Cherbut, G. E. Bergonzelli and S. M. Collins (2010). “Chronic gastrointestinal inflammation induces anxiety-like behavior and alters central nervous system biochemistry in mice.” Gastroenterology 139(6): 2102–2112 e2101.

Brydges, C. R., O. Fiehn, H. S. Mayberg, H. Schreiber, S. M. Dehkordi, S. Bhattacharyya, J. Cha, K. S. Choi, W. E. Craighead, R. R. Krishnan, A. J. Rush, B. W. Dunlop, R. Kaddurah-Daouk and C. Mood Disorders Precision Medicine (2021). “Indoxyl sulfate, a gut microbiome-derived uremic toxin, is associated with psychic anxiety and its functional magnetic resonance imaging-based neurologic signature.” Sci Rep 11(1): 21011.

Caspani, G., S. Kennedy, J. A. Foster and J. Swann (2019). “Gut microbial metabolites in depression: understanding the biochemical mechanisms.” Microb Cell 6(10): 454–481.

Ceryak, S., B. Bouscarel, M. Malavolti and H. Fromm (1998). “Extrahepatic deposition and cytotoxicity of lithocholic acid: studies in two hamster models of hepatic failure and in cultured human fibroblasts.” Hepatology 27(2): 546–556.

Chen, W. G., J. X. Zheng, X. Xu, Y. M. Hu and Y. M. Ma (2018). “Hippocampal FXR plays a role in the pathogenesis of depression: A preliminary study based on lentiviral gene modulation.” Psychiatry Res 264: 374–379.

Chiang, J. Y. L. (2017). “Linking Sex Differences in Non-Alcoholic Fatty Liver Disease to Bile Acid Signaling, Gut Microbiota, and High Fat Diet.” Am J Pathol 187(8): 1658–1659.

Daruich, A., E. Picard, J. H. Boatright and F. Behar-Cohen (2019). “Review: The bile acids urso- and tauroursodeoxycholic acid as neuroprotective therapies in retinal disease.” Mol Vis 25: 610–624.

de Weerth, C. (2017). “Do bacteria shape our development? Crosstalk between intestinal microbiota and HPA axis.” Neurosci Biobehav Rev 83: 458–471.

Devlin, A. S. and M. A. Fischbach (2015). “A biosynthetic pathway for a prominent class of microbiota-derived bile acids.” Nat Chem Biol 11(9): 685–690.

Dinan, T. G. and J. F. Cryan (2015). “The impact of gut microbiota on brain and behaviour: implications for psychiatry.” Curr Opin Clin Nutr Metab Care 18(6): 552–558.

Dinan, T. G. and J. F. Cryan (2017). “Brain-Gut-Microbiota Axis and Mental Health.” Psychosom Med 79(8): 920–926.

Dunlop, B. W., E. B. Binder, J. F. Cubells, M. M. Goodman, M. E. Kelley, B. Kinkead, M. Kutner, C. B. Nemeroff, D. J. Newport, M. J. Owens, T. W. Pace, J. C. Ritchie, V. A. Rivera, D. Westen, W. E. Craighead and H. S. Mayberg (2012). “Predictors of remission in depression to individual and combined treatments (PReDICT): study protocol for a randomized controlled trial.” Trials 13:106.

Dunlop, B. W., M. E. Kelley, V. Aponte-Rivera, T. Mletzko-Crowe, B. Kinkead, J. C. Ritchie, C. B. Nemeroff, W. E. Craighead, H. S. Mayberg and P. R. Team (2017). “Effects of Patient Preferences on Outcomes in the Predictors of Remission in Depression to Individual and Combined Treatments (PReDICT) Study.” Am J Psychiatry 174(6): 546–556.

Dunlop, B. W., D. LoParo, B. Kinkead, T. Mletzko-Crowe, S. P. Cole, C. B. Nemeroff, H. S. Mayberg and W. E. Craighead (2019). “Benefits of Sequentially Adding Cognitive-Behavioral Therapy or Antidepressant Medication for Adults With Nonremitting Depression.” Am J Psychiatry 176(4): 275–286.

Dunlop, B. W., J. K. Rajendra, W. E. Craighead, M. E. Kelley, C. L. McGrath, K. S. Choi, B. Kinkead, C. B. Nemeroff and H. S. Mayberg (2017). “Functional Connectivity of the Subcallosal Cingulate Cortex And Differential Outcomes to Treatment With Cognitive-Behavioral Therapy or Antidepressant Medication for Major Depressive Disorder.” Am J Psychiatry 174(6): 533–545.

Dunlop, B. W., S. Still, D. LoParo, V. Aponte-Rivera, B. N. Johnson, R. L. Schneider, C. B. Nemeroff, H. S. Mayberg and W. E. Craighead (2020). “Somatic symptoms in treatment-naive Hispanic and non-Hispanic patients with major depression.” Depress Anxiety 37(2): 156–165.

Foster, J. A., L. Rinaman and J. F. Cryan (2017). “Stress & the gut-brain axis: Regulation by the microbiome.” Neurobiol Stress 7: 124–136.

Foster, K. R., J. Schluter, K. Z. Coyte and S. Rakoff-Nahoum (2017). “The evolution of the host microbiome as an ecosystem on a leash.” Nature 548(7665): 43–51.

Grant, S. M. and S. DeMorrow (2020). “Bile Acid Signaling in Neurodegenerative and Neurological Disorders.” Int J Mol Sci 21(17).

Guida, F., F. Turco, M. Iannotta, D. De Gregorio, I. Palumbo, G. Sarnelli, A. Furiano, F. Napolitano, S. Boccella, L. Luongo, M. Mazzitelli, A. Usiello, F. De Filippis, F. A. Iannotti, F. Piscitelli, D. Ercolini, V. de Novellis, V. Di Marzo, R. Cuomo and S. Maione (2018). “Antibiotic-induced microbiota perturbation causes gut endocannabinoidome changes, hippocampal neuroglial reorganization and depression in mice.” Brain Behav Immun 67: 230–245.

Hamilton, M. (1959). “The assessment of anxiety states by rating.” Br J Med Psychol 32(1): 50–55.

Hamilton, M. (1960). “A rating scale for depression.” J Neurol Neurosurg Psychiatry 23: 56–62.

Hang, S., D. Paik, L. Yao, E. Kim, J. Trinath, J. Lu, S. Ha, B. N. Nelson, S. P. Kelly, L. Wu, Y. Zheng, R. S. Longman, F. Rastinejad, A. S. Devlin, M. R. Krout, M. A. Fischbach, D. R. Littman and J. R. Huh (2019). “Bile acid metabolites control TH17 and Treg cell differentiation.” Nature 576(7785): 143–148.

Heubi, J. E., K. D. Setchell and K. E. Bove (2007). “Inborn errors of bile acid metabolism.” Semin Liver Dis 27(3): 282–294.

Hibbing, M. E., C. Fuqua, M. R. Parsek and S. B. Peterson (2010). “Bacterial competition: surviving and thriving in the microbial jungle.” Nat Rev Microbiol 8(1): 15–25.

Higashi, T., S. Watanabe, K. Tomaru, W. Yamazaki, K. Yoshizawa, S. Ogawa, H. Nagao, K. Minato, M. Maekawa and N. Mano (2017). “Unconjugated bile acids in rat brain: Analytical method based on LC/ESI-MS/MS with chemical derivatization and estimation of their origin by comparison to serum levels.” Steroids 125:107–113.

Hofmann, A. F. and L. R. Hagey (2008). “Bile acids: chemistry, pathochemistry, biology, pathobiology, and therapeutics.” Cell Mol Life Sci 65(16): 2461–2483.

Huang, F., T. Wang, Y. Lan, L. Yang, W. Pan, Y. Zhu, B. Lv, Y. Wei, H. Shi, H. Wu, B. Zhang, J. Wang, X. Duan, Z. Hu and X. Wu (2015). “Deletion of mouse FXR gene disturbs multiple neurotransmitter systems and alters neurobehavior.” Front Behav Neurosci 9: 70.

Hylemon, P. B., H. Zhou, W. M. Pandak, S. Ren, G. Gil and P. Dent (2009). “Bile acids as regulatory molecules.” J Lipid Res 50(8): 1509–1520.

Kawamata, Y., R. Fujii, M. Hosoya, M. Harada, H. Yoshida, M. Miwa, S. Fukusumi, Y. Habata, T. Itoh, Y. Shintani, S. Hinuma, Y. Fujisawa and M. Fujino (2003). “A G protein-coupled receptor responsive to bile acids.” J Biol Chem 278(11): 9435–9440.

Keller, J., R. Gomez, G. Williams, A. Lembke, L. Lazzeroni, G. M. Murphy, Jr. and A. F. Schatzberg (2017). “HPA axis in major depression: cortisol, clinical symptomatology and genetic variation predict cognition.” Mol Psychiatry 22(4): 527–536.

Kelly, J. R., Y. Borre, O. B. C, E. Patterson, S. El Aidy, J. Deane, P. J. Kennedy, S. Beers, K. Scott, G. Moloney, A. E. Hoban, L. Scott, P. Fitzgerald, P. Ross, C. Stanton, G. Clarke, J. F. Cryan and T. G. Dinan (2016). “Transferring the blues: Depression-associated gut microbiota induces neurobehavioural changes in the rat.” J Psychiatr Res 82:109–118.

Kiriyama, Y. and H. Nochi (2019). “The Biosynthesis, Signaling, and Neurological Functions of Bile Acids.” Biomolecules 9(6).

Kulkarni, M. S., P. M. Heidepriem and K. L. Yielding (1980). “Production by lithocholic acid of DNA strand breaks in L1210 cells.” Cancer Res 40(8 Pt 1): 2666–2669.

Lefebvre, P., B. Cariou, F. Lien, F. Kuipers and B. Staels (2009). “Role of bile acids and bile acid receptors in metabolic regulation.” Physiol Rev 89(1): 147–191.

Li, T. and J. Y. Chiang (2014). “Bile acid signaling in metabolic disease and drug therapy.” Pharmacol Rev 66(4): 948–983.

Li, T. and J. Y. Chiang (2015). “Bile acids as metabolic regulators.” Curr Opin Gastroenterol 31(2): 159–165.

MahmoudianDehkordi, S., M. Arnold, K. Nho, S. Ahmad, W. Jia, G. Xie, G. Louie, A. Kueider-Paisley, M. A. Moseley, J. W. Thompson, L. St John Williams, J. D. Tenenbaum, C. Blach, R. Baillie, X. Han, S. Bhattacharyya, J. B. Toledo, S. Schafferer, S. Klein, T. Koal, S. L. Risacher, M. A. Kling, A. Motsinger-Reif, D. M. Rotroff, J. Jack, T. Hankemeier, D. A. Bennett, P. L. De Jager, J. Q. Trojanowski, L. M. Shaw, M. W. Weiner, P. M. Doraiswamy, C. M. van Duijn, A. J. Saykin, G. Kastenmuller, R. Kaddurah-Daouk, I. Alzheimer’s Disease Neuroimaging and C. the Alzheimer Disease Metabolomics (2019). “Altered bile acid profile associates with cognitive impairment in Alzheimer’s disease-An emerging role for gut microbiome.” Alzheimers Dement 15(1): 76–92.

Marksteiner, J., I. Blasko, G. Kemmler, T. Koal and C. Humpel (2018). “Bile acid quantification of 20 plasma metabolites identifies lithocholic acid as a putative biomarker in Alzheimer’s disease.” Metabolomics 14(1): 1.

Martinot, E., L. Sedes, M. Baptissart, J. M. Lobaccaro, F. Caira, C. Beaudoin and D. H. Volle (2017). “Bile acids and their receptors.” Mol Aspects Med 56: 2–9.

Matza, L. S., R. Morlock, C. Sexton, K. Malley and D. Feltner (2010). “Identifying HAM-A cutoffs for mild, moderate, and severe generalized anxiety disorder.” Int J Methods Psychiatr Res 19(4): 223–232.

Milaneschi, Y., M. Arnold, G. Kastenmuller, S. M. Dehkordi, R. R. Krishnan, B. W. Dunlop, A. J. Rush, B. W. Penninx and R. Kaddurah-Daouk (2021). “Genomics-based identification of a potential causal role for acylcarnitine metabolism in depression.” medRxiv.

Miranda, M., J. F. Morici, M. B. Zanoni and P. Bekinschtein (2019). “Brain-Derived Neurotrophic Factor: A Key Molecule for Memory in the Healthy and the Pathological Brain.” Front Cell Neurosci 13: 363.

Monteiro-Cardoso, V. F., M. Corliano and R. R. Singaraja (2021). “Bile Acids: A Communication Channel in the Gut-Brain Axis.” Neuromolecular Med 23(1): 99–117.

Mortiboys, H., J. Aasly and O. Bandmann (2013). “Ursocholanic acid rescues mitochondrial function in common forms of familial Parkinson’s disease.” Brain 136(Pt 10): 3038–3050.

Nho, K., A. Kueider-Paisley, S. MahmoudianDehkordi, M. Arnold, S. L. Risacher, G. Louie, C. Blach, R. Baillie, X. Han, G. Kastenmuller, W. Jia, G. Xie, S. Ahmad, T. Hankemeier, C. M. van Duijn, J. Q. Trojanowski, L. M. Shaw, M. W. Weiner, P. M. Doraiswamy, A. J. Saykin, R. Kaddurah-Daouk, I. Alzheimer’s Disease Neuroimaging and C. the Alzheimer Disease Metabolomics (2019). “Altered bile acid profile in mild cognitive impairment and Alzheimer’s disease: Relationship to neuroimaging and CSF biomarkers.” Alzheimers Dement 15(2): 232–244.

Nie, S., G. Chen, X. Cao and Y. Zhang (2014). “Cerebrotendinous xanthomatosis: a comprehensive review of pathogenesis, clinical manifestations, diagnosis, and management.” Orphanet J Rare Dis 9:179.

O’Byrne, J., M. C. Hunt, D. K. Rai, M. Saeki and S. E. Alexson (2003). “The human bile acid-CoA:amino acid N-acyltransferase functions in the conjugation of fatty acids to glycine.” J Biol Chem 278(36): 34237–34244.

O’Sullivan, E., E. Barrett, S. Grenham, P. Fitzgerald, C. Stanton, R. P. Ross, E. M. Quigley, J. F. Cryan and T. G. Dinan (2011). “BDNF expression in the hippocampus of maternally separated rats: does Bifidobacterium breve 6330 alter BDNF levels?” Benef Microbes 2(3): 199–207.

Odermatt, A., T. Da Cunha, C. A. Penno, C. Chandsawangbhuwana, C. Reichert, A. Wolf, M. Dong and M. E. Baker (2011). “Hepatic reduction of the secondary bile acid 7-oxolithocholic acid is mediated by 11beta-hydroxysteroid dehydrogenase 1.” Biochem J 436(3): 621–629.

Palmer, R. H. and M. G. Bolt (1971). “Bile acid sulfates. I. Synthesis of lithocholic acid sulfates and their identification in human bile.” J Lipid Res 12(6): 671–679.

Parada Venegas, D., M. K. De la Fuente, G. Landskron, M. J. Gonzalez, R. Quera, G. Dijkstra, H. J. M. Harmsen, K. N. Faber and M. A. Hermoso (2019). “Short Chain Fatty Acids (SCFAs)-Mediated Gut Epithelial and Immune Regulation and Its Relevance for Inflammatory Bowel Diseases.” Front Immunol 10: 277.

Parry, G. J., C. M. Rodrigues, M. M. Aranha, S. J. Hilbert, C. Davey, P. Kelkar, W. C. Low and C. J. Steer (2010). “Safety, tolerability, and cerebrospinal fluid penetration of ursodeoxycholic Acid in patients with amyotrophic lateral sclerosis.” Clin Neuropharmacol 33(1): 17–21.

Pauli-Magnus, C. and P. J. Meier (2005). “Hepatocellular transporters and cholestasis.” J Clin Gastroenterol 39(4 Suppl 2): S103–110.

Qiu, Y., G. Cai, M. Su, T. Chen, X. Zheng, Y. Xu, Y. Ni, A. Zhao, L. X. Xu, S. Cai and W. Jia (2009). “Serum metabolite profiling of human colorectal cancer using GC-TOFMS and UPLC-QTOFMS.” J Proteome Res 8(10): 4844–4850.

Ramalho, R. M., A. F. Nunes, R. B. Dias, J. D. Amaral, A. C. Lo, R. D’Hooge, A. M. Sebastiao and C. M. Rodrigues (2013). “Tauroursodeoxycholic acid suppresses amyloid beta-induced synaptic toxicity in vitro and in APP/PS1 mice.” Neurobiol Aging 34(2): 551–561.

Ridlon, J. M. and J. S. Bajaj (2015). “The human gut sterolbiome: bile acid-microbiome endocrine aspects and therapeutics.” Acta Pharm Sin B 5(2): 99–105.

Ridlon, J. M., S. C. Harris, S. Bhowmik, D. J. Kang and P. B. Hylemon (2016). “Consequences of bile salt biotransformations by intestinal bacteria.” Gut Microbes 7(1): 22–39.

Rieder, R., P. J. Wisniewski, B. L. Alderman and S. C. Campbell (2017). “Microbes and mental health: A review.” Brain Behav Immun 66: 9–17.

Rodrigues, C. M., C. L. Stieers, C. D. Keene, X. Ma, B. T. Kren, W. C. Low and C. J. Steer (2000). “Tauroursodeoxycholic acid partially prevents apoptosis induced by 3-nitropropionic acid: evidence for a mitochondrial pathway independent of the permeability transition.” J Neurochem 75(6): 2368–2379.

Shonsey, E. M., M. Sfakianos, M. Johnson, D. He, C. N. Falany, J. Falany, D. J. Merkler and S. Barnes (2005). “Bile acid coenzyme A: amino acid N-acyltransferase in the amino acid conjugation of bile acids.” Methods Enzymol 400: 374–394.

Simpson, C. A., C. Diaz-Arteche, D. Eliby, O. S. Schwartz, J. G. Simmons and C. S. M. Cowan (2021). “The gut microbiota in anxiety and depression - A systematic review.” Clin Psychol Rev 83: 101943.

Simpson, C. A., O. S. Schwartz and J. G. Simmons (2020). “The human gut microbiota and depression: widely reviewed, yet poorly understood.” J Affect Disord 274: 73–75.

Sonne, D. P., M. Hansen and F. K. Knop (2014). “Bile acid sequestrants in type 2 diabetes: potential effects on GLP1 secretion.” Eur J Endocrinol 171(2): R47–65.

Sudo, N., Y. Chida, Y. Aiba, J. Sonoda, N. Oyama, X. N. Yu, C. Kubo and Y. Koga (2004). “Postnatal microbial colonization programs the hypothalamic-pituitary-adrenal system for stress response in mice.” J Physiol 558(Pt 1): 263–275.

Sundaram, S. S., K. E. Bove, M. A. Lovell and R. J. Sokol (2008). “Mechanisms of disease: Inborn errors of bile acid synthesis.” Nat Clin Pract Gastroenterol Hepatol 5(8): 456–468.

Tognini, P. (2017). “Gut Microbiota: A Potential Regulator of Neurodevelopment.” Front Cell Neurosci 11:25.

Tremlett, H., K. C. Bauer, S. Appel-Cresswell, B. B. Finlay and E. Waubant (2017). “The gut microbiome in human neurological disease: A review.” Ann Neurol 81(3): 369–382.

Vaz, F. M. and S. Ferdinandusse (2017). “Bile acid analysis in human disorders of bile acid biosynthesis.” Mol Aspects Med 56:10–24.

Wahlstrom, A., S. I. Sayin, H. U. Marschall and F. Backhed (2016). “Intestinal Crosstalk between Bile Acids and Microbiota and Its Impact on Host Metabolism.” Cell Metab 24(1): 41–50.

Wan, Y. Y. and L. Sheng (2018). “Regulation of bile acid receptor activity().” Liver Res 2(4): 180–185.

Weitz, E. S., S. D. Hollon, J. Twisk, A. van Straten, M. J. Huibers, D. David, R. J. DeRubeis, S. Dimidjian, B. W. Dunlop, I. A. Cristea, M. Faramarzi, U. Hegerl, R. B. Jarrett, F. Kheirkhah, S. H. Kennedy, R. Mergl, J. Miranda, D. C. Mohr, A. J. Rush, Z. V. Segal, J. Siddique, A. D. Simons, J. R. Vittengl and P. Cuijpers (2015). “Baseline Depression Severity as Moderator of Depression Outcomes Between Cognitive Behavioral Therapy vs Pharmacotherapy: An Individual Patient Data Meta-analysis.” JAMA Psychiatry 72(11): 1102–1109.

Xie, G., X. Wang, R. Jiang, A. Zhao, J. Yan, X. Zheng, F. Huang, X. Liu, J. Panee, C. Rajani, C. Yao, H. Yu, W. Jia, B. Sun, P. Liu and W. Jia (2018). “Dysregulated bile acid signaling contributes to the neurological impairment in murine models of acute and chronic liver failure.” EBioMedicine 37: 294–306.

Xie, G., Y. Wang, X. Wang, A. Zhao, T. Chen, Y. Ni, L. Wong, H. Zhang, J. Zhang, C. Liu, P. Liu and W. Jia (2015). “Profiling of serum bile acids in a healthy Chinese population using UPLC-MS/MS.” J Proteome Res 14(2): 850–859.

Yarandi, S. S., D. A. Peterson, G. J. Treisman, T. H. Moran and P. J. Pasricha (2016). “Modulatory Effects of Gut Microbiota on the Central Nervous System: How Gut Could Play a Role in Neuropsychiatric Health and Diseases.” J Neurogastroenterol Motil 22(2): 201–212.

Zhao, L., Y. Ni, M. Su, H. Li, F. Dong, W. Chen, R. Wei, L. Zhang, S. P. Guiraud, F. P. Martin, C. Rajani, G. Xie and W. Jia (2017). “High Throughput and Quantitative Measurement of Microbial Metabolome by Gas Chromatography/Mass Spectrometry Using Automated Alkyl Chloroformate Derivatization.” Anal Chem 89(10): 5565–5577.

Zheng, P., B. Zeng, C. Zhou, M. Liu, Z. Fang, X. Xu, L. Zeng, J. Chen, S. Fan, X. Du, X. Zhang, D. Yang, Y. Yang, H. Meng, W. Li, N. D. Melgiri, J. Licinio, H. Wei and P. Xie (2016). “Gut microbiome remodeling induces depressive-like behaviors through a pathway mediated by the host’s metabolism.” Mol Psychiatry 21(6): 786–796.

