## Supplemental Results for "Gut microbiome-linked metabolites in the pathobiology of depression and anxiety - a role for bile acids"

**Supplementary Methods**

***Partial Least-Squares Regression and Discriminant Analysis:***

We conducted separate partial least squares regression (PLS-regression) and discriminant analysis (PLSDA) to examine contribution of baseline BA levels to baseline HRSD_17_, HRSA-total and treatment outcome. In all models, we accounted for age, sex and BMI and used 5-fold cross-validation with 100 repeats. In PLS-regression models, baseline BA profiles of all patients were considered as predictor variables and HRSD_17_ and HRSA total as continuous dependent variables. Using a PLSDA model, we examined if the baseline BA profiles could discriminate patients at the two extremes of the treatment response spectrum, the remitters, and the treatment failures. Significant predictors were identified based on their variable importance on projection (VIP) scores. Variables with a VIP value >1 were considered important for the models.

**PLS Analyses RESULTS:**

Variable importance projection (VIP) scores from PLS analyses are presented in **Supplementary Table 3.** PLS regression model with HRSD_17_ as the dependent variable resulted in 6 significant BAs (VIP score > 1), 4 primary (including both conjugated and non-conjugated) and 2 secondary BAs as important contributors to depression severity. These were GLCA_3S (VIP=3.12), CDCA (VIP=2.95), GCDCA (VIP=2.84), GCA(VIP=1.86), TCA(VIP=1.31), and UDCA(VIP=1.21)).

PLS regression model with total HRSA score as the dependent variable resulted in 3 primary and 2 secondary BAs, GLCA_3s (VIP=3.28), GDCA (VIP=3.06), GCDCA (VIP=3.05), GCA (VIP=1.28), CDCA (VIP=1.24) and CDCA_24G (VIP=1.11) as important contributors to anxiety symptoms.

A partial least squares discriminant analysis (PLSDA) was also conducted to investigate which bile acids could most accurately distinguish between remitters and failures at baseline. The first component of the PLSDA model achieved an area under the curve of 0.7923 (*p* = 0.00002). Based on the VIP scores, CDCA (more abundant in remitters) was the most important variable in discriminating remitters from treatment failures.

Supplementary data:

**Supplemental Table 1**- Serum levels of bile acids primary, secondary and their conjugated forms in patients enrolled in PReDICT study. All values are at baseline prior to treatment.

| **Metabolite** | **Abbreviate** | **HMDB ID** | **Type** | **Conjugation?** | **Population (N=208)** | **Remission (N=73)** | **Treatment Failure (N=25)** |
| --- | --- | --- | --- | --- | --- | --- | --- |
| Cholic Acid | CA | HMDB00619 | Primary | Unconjugated | 268.79(26.61) | 223.02(25.21) | 189.76(32.02) |
| Chenodeoxycholic Acid | CDCA | HMDB00518 | Primary | Unconjugated | 417.81(36.10) | 433.58(47.12) | 200.08(23.28) |
| Chenodeoxycholic Acid 24 Acyl β D glucuronide | CDCA_24G | NA | Primary | Unconjugated | 285.92(21.28) | 331.33(36.98) | 257.71(53.79) |
| 3β Cholic Acid | βCA | HMDB00419 | Primary | Unconjugated | 26.88(1.03) | 27.75(1.83) | 22.85(2.06) |
| Hyocholic Acid | HCA | HMDB00760 | Primary | Unconjugated | 137.82(44.29) | 152.39(72.50) | 53.57(27.53) |
| ω Muricholic Acid | ωMCA | HMDB00364 | Primary | Unconjugated | 70.65(2.29) | 64.87(3.08) | 62.97(5.64) |
| Nor Cholic acid | NorCA | NA | Primary | Unconjugated | 26.03(2.01) | 24.38(2.97) | 23.71(5.47) |
| Taurocholic Acid | TCA | HMDB00036 | Primary | Conjugated | 119.14(18.02) | 109.20(14.50) | 121.85(40.14) |
| Glycocholic Acid | GCA | HMDB00138 | Primary | Conjugated | 485.04(45.56) | 522.85(68.65) | 388.60(88.50) |
| Taurochenodeoxycholate | TCDCA | HMDB00951 | Primary | Conjugated | 245.74(26.16) | 229.21(29.42) | 201.18(48.28) |
| Glycochenodeoxycholate | GCDCA | HMDB00637 | Primary | Conjugated | 1578.69(104.72) | 1775.25(168.81) | 1291.23(210.95) |
| 5β cholanic acid 3β,7β,12α triol 5β Cholanic Acid 3β, 7β, 12α triol | βUCA | M07X096_N | Primary | Unconjugated | 25.03(1.66) | 24.12(2.66) | 22.54(3.48) |
| Taurohyocholate | THCA | HMDB11637 | Primary | Conjugated | 52.68(4.72) | 53.51(8.65) | 34.80(6.56) |
| Glycohyocholate | GHCA | M07X072_N | Primary | Conjugated | 7.37(0.37) | 6.68(0.49) | 9.72(1.37) |
| Glycohyodeoxycholate | GHDCA | NA | Secondary | Conjugated | 6.42(0.40) | 6.53(0.62) | 5.41(0.92) |
| Deoxycholic Acid | DCA | HMDB00626 | Secondary | Unconjugated | 643.60(41.72) | 568.73(59.21) | 469.11(66.41) |
| 3Œ±,12Œ± Dihydroxynorcholanate\23 Nordeoxycholic acid | NorDCA | NA | Secondary | Unconjugated | 12.29(0.81) | 12.77(1.10) | 13.19(4.12) |
| Ursodeoxycholic Acid | UDCA | HMDB00946 | Secondary | Unconjugated | 124.34(14.83) | 163.84(39.10) | 82.60(14.42) |
| Lithocholic Acid | LCA | HMDB00761 | Secondary | Unconjugated | 21.51(1.74) | 18.22(1.61) | 17.09(2.73) |
| 6,7 diketolithocholic acid | 6,7_diketoLCA | NA | Secondary | Unconjugated | 159.38(10.43) | 153.24(16.20) | 141.61(32.55) |
| Nutriacholic acid / 7 Ketolithocholic Acid | 7_ketoLCA | HMDB00467 | Secondary | Unconjugated | 12.79(3.21) | 6.79(0.67) | 30.21(23.90) |
| Lithocholic Acid 3 Sulfate | LCA_3S | HMDB00907 | Secondary | Unconjugated | 21.86(0.91) | 19.51(1.02) | 19.31(2.29) |
| Hyodeoxycholic Acid | HDCA | HMDB00733 | Secondary | Unconjugated | 92.22(7.96) | 97.19(14.07) | 65.02(13.37) |
| Isolithocholic acid | isoLCA | HMDB00717 | Secondary | Unconjugated | 38.32(2.58) | 41.33(5.24) | 22.98(5.87) |
| βHyodeoxycholic Acid (Isohyodeoxycholic acid) | βHDCA | HMDB00664 | Secondary | Unconjugated | 25.30(2.62) | 22.60(4.36) | 21.25(4.89) |
| Allolithocholic acid ( Isoallolithocholic acid) | alloLCA | HMDB00713 | Secondary | Unconjugated | 36.49(2.42) | 33.85(4.01) | 27.46(5.97) |
| dehydroLCA | dehydroLCA | M07X058a_N2 | Secondary | Unconjugated | 14.14(0.47) | 14.45(0.73) | 13.75(0.92) |
| 12 Ketolithocholic acid\12 Ketodeoxycholic acid | 12_ketoLCA | HMDB00328 | Secondary | Unconjugated | 7.45(0.51) | 6.18(0.59) | 4.91(0.81) |
| 3β Ursodeoxycholic Acid (Isoursodeoxycholic acid) | 3β UDCA | HMDB00686 | Secondary | Unconjugated | 128.07(13.22) | 157.13(29.53) | 90.31(16.49) |
| Glycodeoxycholic Acid | GDCA | HMDB00631 | Secondary | Conjugated | 707.07(45.64) | 667.53(71.78) | 549.05(100.32) |
| Tauroursodeoxycholic Acid | TUDCA | HMDB00874 | Secondary | Conjugated | 20.42(1.21) | 22.85(2.13) | 24.49(5.12) |
| Glycoursodeoxycholic Acid | GUDCA | HMDB00708 | Secondary | Conjugated | 13.86(1.01) | 14.79(1.97) | 11.12(2.86) |
| Taurodeoxycholic Acid | TDCA | HMDB00896 | Secondary | Conjugated | 71.37(7.21) | 71.81(9.45) | 53.42(8.54) |
| Glycolithocholic Acid 3 Sulfate | GLCA_3S | HMDB02639 | Secondary | Conjugated | 800.54(74.44) | 650.96(73.37) | 788.42(219.88) |
| Glycolithocholate | GLCA | HMDB00698 | Secondary | Conjugated | 5.08(0.34) | 5.03(0.65) | 4.53(0.90) |
| Taurohyodeoxycholic Acid | THDCA | M07X076_N | Secondary | Conjugated | 15.66(1.73) | 11.86(1.59) | 11.17(2.08) |

**Supplementary Table 2**: BA ratios, summations, and their association with metabolic reactions and dysfunction.

| **Ratios indicating metabolic reactions in BA pathway** | **Formula** | **Description** | **Findings in PReDICT** | **Literature** |
| --- | --- | --- | --- | --- |
| 7a-Dehydroxylation of CA | DCA / CA | Conversion of the primary bile acid CA to the secondary bile acid DCA by gut bacteria. The ratio is strongly associated with a cognitive decline. | N.S | MahmoudianDehkordi et al. 2019, PMID: 30337151. |
| GDCA Synthesis from CA | GDCA / CA | Conversion of the primary bile acid CA to the conjugated secondary bile acid GDCA involving the gut microbiota. The ratio is strongly associated with cognitive decline. | Higher ratio values correlates with higher HRSA-Total scores | MahmoudianDehkordi et al. 2019, PMID: 30337151. |
| GLCA Synthesis from CDCA | GLCA / CDCA | Conversion of the primary bile acid CDCA to the conjugated secondary bile acid GLCA involving the gut microbiota. The ratio is strongly associated with cognitive decline. | Higher ratio values correlates with higher HRSA-Total scores | MahmoudianDehkordi et al. 2019, PMID: 30337151. |
| Gly Conjugation of CA | GCA / CA | Indicator of conjugation of glycine to the primary bile acid CA to form GCA. | N.S | Shonsey et al. 2005, DOI: 10.1016/S0076-6879(05)00022-4. |
| Gly Conjugation of CDCA | GCDCA / CDCA | Indicator of conjugation of glycine to the primary bile acid CDCA to form GCDCA. | N.S | Shonsey et al. 2005, DOI: 10.1016/S0076-6879(05)00022-4. |
| Gly Conjugation of DCA | GDCA / DCA | Indicator of conjugation of glycine to the secondary bile acid DCA to form GDCA. | Higher ratio values correlates with higher HRSA-Total scores | Shonsey et al. 2005, DOI: 10.1016/S0076-6879(05)00022-4. |
| Gly Conjugation of Primary BAs | (GCA + GCDCA) / (CA + CDCA) | Indicator of conjugation of glycine to the primary bile acids CA and CDCA to form the glycine-conjugated primary bile acids GCA and GCDCA. | N.S | Shonsey et al. 2005, DOI: 10.1016/S0076-6879(05)00022-4. |
| Primary BA Conjugation | (GCA + GCDCA + TCA + TCDCA) / (CA + CDCA) | Indicator of conjugation of glycine or taurine to primary bile acids to form conjugated primary bile acids in the liver. | N.S | Shonsey et al. 2005, DOI: 10.1016/S0076-6879(05)00022-4;  O'Byrne et al. 2003, DOI: 10.1074/jbc.M300987200. |
| Ratio of 12a-OH BAs to Non-12a-OH BAs | (CA + DCA + GCA + GDCA + TCA +  TDCA) / (CDCA + GCDCA + GLCA +  GUDCA + TCDCA + TLCA) | The ratio of 12-alpha-hydroxylated bile acids to non-12-alpha-hydroxylated bile acids is an indicator of type 2 diabetes. | Higher ratio values correlates with higher HRSA-Total scores | Chiang 2017, DOI: 10.1016/j.ajpath.2017.06.001;  Sonne et al. 2014, DOI: 10.1530/eje-14-0154. |
| Ratio of CDCA to CA | CDCA / CA | Ratio of the two primary bile acids. CA is synthesized from cholesterol in the classical pathway, while CDCA primarily comes from the alternative pathway. | N.S | Vaz and Ferdinandusse 2010, PMID: 28322867;  Huang et al. 2009, DOI: 10.1055/s-0028-1103158. |
| Ratio of Primary BAs to BAs | (CA + CDCA + GCA + GCDCA + TCA + TCDCA) / (CA + CDCA + DCA + GCA + GCDCA + GDCA + GLCA + GUDCA +  TCA + TCDCA + TDCA + TLCA) | Fraction of primary bile acids (BAs) relative to the BA pool. Primary BAs are synthesized from cholesterol in the liver, conjugated with either taurine or glycine, and then released into the biliary system. After their excretion into the gastrointestinal tract, some of them are deconjugated and modified by bacterial metabolism, leading to secondary BAs. | Higher ratio values correlates with lower HRSA-Total scores | Boyer 2013, DOI: 10.1002/cphy.c120027. |
| Ratio of Secondary BAs to BAs | (DCA + GDCA + GLCA + GUDCA +  TDCA + TLCA) / (CA + CDCA + DCA +  GCA + GCDCA + GDCA + GLCA +  GUDCA + TCA + TCDCA + TDCA + TLCA) | Fraction of secondary bile acids (BAs) relative to the BA pool. | Higher ratio values correlates with higher HRSA-Total scores | Boyer 2013, DOI: 10.1002/cphy.c120027. |
| Secondary BA Conjugation | (GDCA + GLCA + GUDCA + TDCA +  TLCA) / DCA | Indicator of conjugation of glycine or taurine to secondary bile acids to form conjugated secondary bile acids in the liver. | Higher ratio values weakly correlates with higher HRSA-Total scores | Shonsey et al. 2005, DOI: 10.1016/S0076-6879(05)00022-4;  O'Byrne et al. 2003, DOI: 10.1074/jbc.M300987200. |
| Secondary BA Synthesis | (DCA + GDCA + GLCA + GUDCA +  TDCA + TLCA) / (CA + CDCA + GCA + GCDCA + TCA + TCDCA) | Ratio of cytotoxic secondary bile acids (BAs) to primary BAs. Secondary BAs solely produced by intestinal bacteria can accumulate to a high degree in the enterohepatic circulation of some individuals and may contribute to the pathogenesis of colon cancer, gallstones and other gastrointestinal diseases. | Higher ratio values correlates with higher HRSA-Total scores | Nho et al. 2019, DOI: 10.1016/j.jalz.2018.08.012;  Marksteiner et al. 2017, DOI: 10.1007/s11306-017-1297-5;  Mouzaki et al. 2016, DOI: 10.1371/journal.pone.0151829. |
| Sum of 12a-OH BAs | CA + DCA + GCA + GDCA + TCA + TDCA | It has been shown that increased levels of 12-alpha-hydroxylated bile acids are associated with insulin resistance. | N.S | Chiang 2017, DOI: 10.1016/j.ajpath.2017.06.001;  Tomkin and Owens 2016, DOI: 10.1515/jtim-2016-0018;  Sonne et al. 2014, DOI: 10.1530/eje-14-0154. |
| Sum of BAs | CA + CDCA + DCA + GCA + GCDCA + GDCA + GLCA + GUDCA + TCA +  TCDCA + TDCA + TLCA | Sum of BAs is considered to be a biomarker of liver function. | N.S | Luo et al. 2018, DOI: 10.1371/journal.pone.0193824;  Li and Chiang 2015, DOI: 10.1097/mog.0000000000000156;  Sugita et al. 2015, DOI: 10.1155/2015/717431. |
| Sum of Conjugated BAs | GCA + GCDCA + GDCA + GLCA +  GUDCA + TCA + TCDCA + TDCA + TLCA | Conjugation with Gly or Tau increase solubility of the BAs and make them impermeable to cell membranes. Conjugated BAs may be increased in plasma due to mutations or antibiotic treatments. | N.S | Li et al. 2017, doi: 10.1093/toxsci/kfx015;  Vaz et al. 2015, doi: 10.1002/hep.27240;  Shonsey et al. 2005, DOI: 10.1016/S0076-6879(05)00022-4;  Hofmann 1999, DOI:10.1001/archinte.159.22.2647. |
| Sum of Conjugated Primary BAs | GCA + GCDCA + TCA + TCDCA | Conjugated primary BAs are significantly elevated in patients with polycystic ovary syndrome. In addition, conjugated primary BAs were found to be elevated after antibiotic treatment. | N.S | Zhang et al. 2019; DOI: 10.1016/j.jsbmb.2019.03.005;  Behr et al. 2019, DOI: 10.1016/j.taap.2018.11.012;  Shonsey et al. 2005, DOI: 10.1016/S0076-6879(05)00022-4. |
| Sum of Conjugated Secondary BAs | GDCA + GLCA + GUDCA + TDCA + TLCA | Concentrations of conjugated secondary BAs may be reduced after antibiotic treatment. | Higher ratio values correlates with higher HRSA-Total scores | Behr et al. 2019, DOI: 10.1016/j.taap.2018.11.012;  Shonsey et al. 2005, DOI: 10.1016/S0076-6879(05)00022-4. |
| Sum of Gly-Conjugated BAs | GCA + GCDCA + GDCA + GLCA + GUDCA | The content of taurine-conjugated BAs normally correlates with the content of glycine-conjugated BAs, unless the levels of taurine or glycine are abnormal. | N.S | Li et al. 2017, doi: 10.1093/toxsci/kfx015;  Vaz et al. 2015, doi: 10.1002/hep.27240;  Shonsey et al. 2005, DOI: 10.1016/S0076-6879(05)00022-4;  Hofmann 1999, DOI:10.1001/archinte.159.22.2647. |
| Sum of Non-12a-OH BAs | CDCA + GCDCA + GLCA + GUDCA + TCDCA + TLCA | Sum of non-12-alpha-hydroxylated bile acids. | N.S | Chiang 2017, DOI: 10.1016/j.ajpath.2017.06.001;  Sonne et al. 2014, DOI: 10.1530/eje-14-0154. |
| Sum of Primary BAs | CA + CDCA + GCA + GCDCA + TCA + TCDCA | Elevated primary BA levels in plasma or serum are often caused by biliary obstruction, e.g. in cholelithiasis (gallstones), pancreatitis, or pancreatic cancer. | N.S | Martinot et al. 2017, DOI: 10.1016/j.mam.2017.01.006;  Wahlstroem et al. 2016, DOI: 10.1016/j.cmet.2016.05.005;  Lerch et al. 2010, DOI:10.1053/j.gastro.2009.12.012. |
| Sum of Secondary BAs | DCA + GDCA + GLCA + GUDCA +  TDCA + TLCA | High fat and high beef diets promote bile discharge and influence the bacterial composition in the gut, resulting in increased levels of secondary BAs associated with an increased risk for colon cancer. Low levels of secondary BAs may indicate a reduced capacity of the gut microbiota to metabolize primary BAs. | Higher ratio values correlates with higher HRSA-Total scores | Wahlstroem et al. 2019, DOI: 10.1016/j.cmet.2016.05.005;  Martinot et al. 2017, DOI: 10.1016/j.mam.2017.01.006;  Wahlstroem et al. 2016, DOI: 10.1016/j.cmet.2016.05.005;  Ajouz et al. 2014, DOI: 10.1186/1477-7819-12-164;  Bayerdoerffer et al. 1995, DOI: 10.1136/gut.36.2.268. |
| Sum of Taurine-Conjugated BAs | TCA + TCDCA + TDCA + TLCA | The content of taurine-conjugated BAs normally correlates with the content of glycine-conjugated BAs, unless the levels of taurine or glycine are abnormal. | N.S | Li et al. 2017, DOI: 10.1093/toxsci/kfx015;  Vaz et al. 2015, DOI: 10.1002/hep.27240;  Shonsey et al. 2005, DOI: 10.1016/S0076-6879(05)00022-4;  Hofmann 1999, DOI:10.1001/archinte.159.22.2647. |
| Sum of Unconjugated BAs | CA + CDCA + DCA | Bile acids (BAs) are synthesized in the liver and then usually conjugated with taurine or glycine to increase solubility and make them impermeable to cell membranes, only a fraction of BAs remains unconjugated. Unconjugated BAs may be increased in several hepatobiliary diseases, e.g. cholangiocarcinoma and hepatocellular carcinoma. | N.S | Li et al. 2017, DOI: 10.1016/j.livres.2017.04.001;  Chiang et al. 2017, DOI: 10.12688/f1000research.12449.1;  Shonsey et al. 2005, DOI: 10.1016/S0076-6879(05)00022-4; Changbumrung et al. 1990, PMID: 2161896;  Makino et al. 1969, PMID: 5788911. |
| Sum of Unconjugated Primary BAs | CA + CDCA | Unconjugated primary bile acids (BAs) are produced from cholesterol in the liver and are less hydrophilic than upon conjugation to glycine or taurine to increase solubility. Elevated primary BA levels in plasma or serum are often caused by biliary obstruction, e.g. in cholelithiasis (gallstones), pancreatitis, or pancreatic cancer. | Higher ratio values correlates with lower HRSA-Total scores | Fan et al. 2018, DOI: 10.1515/hsz-2018-0379; Dawson et al. 2015, DOI: 10.1194/jlr.R054114. |
| Taurine Conjugation of CA | TCA / CA | The ratio of the taurine-conjugated primary bile acid taurocholic acid to the unconjugated primary bile acid cholic acid is an indicator of bile acid CoA ligase and bile acid CoA:amino acid N-acyltransferase activity in the liver. | Higher ratio values correlates with higher HRSA-SOM scores | Shonsey et al. 2005, DOI: 10.1016/S0076-6879(05)00022-4. |
| Taurine Conjugation of CDCA | TCDCA / CDCA | The ratio of the taurine-conjugated primary bile acid taurochenodeoxycholic acid to the unconjugated primary bile acid chenodeoxycholic acid is an indicator of bile acid CoA ligase and bile acid CoA:amino acid N-acyltransferase activity in the liver. | N.S | Shonsey et al. 2005, DOI: 10.1016/S0076-6879(05)00022-4. |
| Taurine Conjugation of DCA | TDCA / DCA | The ratio of the taurine-conjugated secondary bile acid taurodeoxycholic acid to the unconjugated secondary bile acid deoxycholic acid is an indicator of bile acid CoA ligase and bile acid CoA:amino acid N-acyltransferase activity in the liver. | N.S | Shonsey et al. 2005, DOI: 10.1016/S0076-6879(05)00022-4. |
| Taurine Conjugation of Primary BAs | (TCA + TCDCA) / (CA + CDCA) | The ratio of the taurine-conjugated primary bile acids taurocholic acid and taurochenodeoxycholic acid to the unconjugated primary bile acids cholic acid and chenodeoxycholic acid is an indicator of bile acid CoA ligase and bile acid CoA:amino acid N-acyltransferase activity in the liver. | N.S | Shonsey et al. 2005, DOI: 10.1016/S0076-6879(05)00022-4. |
| TDCA Synthesis from CA | TDCA / CA | Ratio of the taurine-conjugated secondary bile acid taurodeoxycholic acid to the unconjugated primary bile acid cholic acid, which is strongly associated with cognitive decline. | N.S | MahmoudianDehkordi et al. 2019, PMID: 30337151. |

**Supplementary Table 3:** PLSDA scores for contribution of BA to severity of clinical symptoms.

| **Metabolite** | **Anxiety VIP Score** | **Depression VIP Score** | **Treatment Outcome VIP Score** |
| --- | --- | --- | --- |
| GLCA_3S | 3.28 | 3.12 | 0.49 |
| GDCA | 3.06 | 0.95 | 0.94 |
| GCDCA | 3.05 | 2.84 | 1.35 |
| GCA | 1.28 | 1.86 | 0.93 |
| CDCA | 1.24 | 2.95 | 2.45 |
| CDCA_24G | 1.11 | 0 | 0.87 |
| TCA | 0.74 | 1.31 | 0.6 |
| CA | 0.64 | 0.26 | 0.81 |
| TCDCA | 0.5 | 0.66 | 0.18 |
| 6,7_diketoLCA | 0.45 | 0.32 | 0.17 |
| UDCA | 0.38 | 1.21 | 1.05 |
| DCA | 0.36 | 0.76 | 1.19 |
| HCA | 0.36 | 0.52 | 0.8 |
| βHDCA | 0.32 | 0.29 | 0.37 |
| TDCA | 0.22 | 0.22 | 1.01 |
| isoLCA | 0.18 | 0.14 | 1.71 |
| THCA | 0.17 | 0.13 | 1.24 |
| NorCA | 0.17 | 0.12 | 0.15 |
| βUDCA | 0.15 | 0.55 | 1.21 |
| alloLCA | 0.12 | 0.03 | 0.68 |
| ωMCA | 0.11 | 0.14 | 0.38 |
| βUCA | 0.08 | 0.1 | 0.12 |
| 7_ketoLCA | 0.07 | 0.08 | 1.37 |
| HDCA | 0.06 | 0.14 | 1.17 |
| LCA_3S | 0.06 | 0.06 | 0.1 |
| 12_ketoLCA | 0.04 | 0.02 | 1.2 |
| GUDCA | 0.04 | 0.04 | 0.93 |
| LCA | 0.04 | 0.02 | 0.22 |
| GHDCA | 0.03 | 0.02 | 0.62 |
| dehydroLCA | 0.03 | 0.03 | 0.53 |
| TUDCA | 0.03 | 0.04 | 0.42 |
| THDCA | 0.03 | 0 | 0.28 |
| NorDCA | 0.03 | 0.01 | 0.03 |
| GHCA | 0.01 | 0.02 | 2.19 |
| βCA | 0.01 | 0.06 | 1.45 |
| GLCA | 0 | 0.02 | 0.14 |

Supplementary Figure 1: Correlation of baseline clinical symptoms ofn Emory PReDICT Cohort

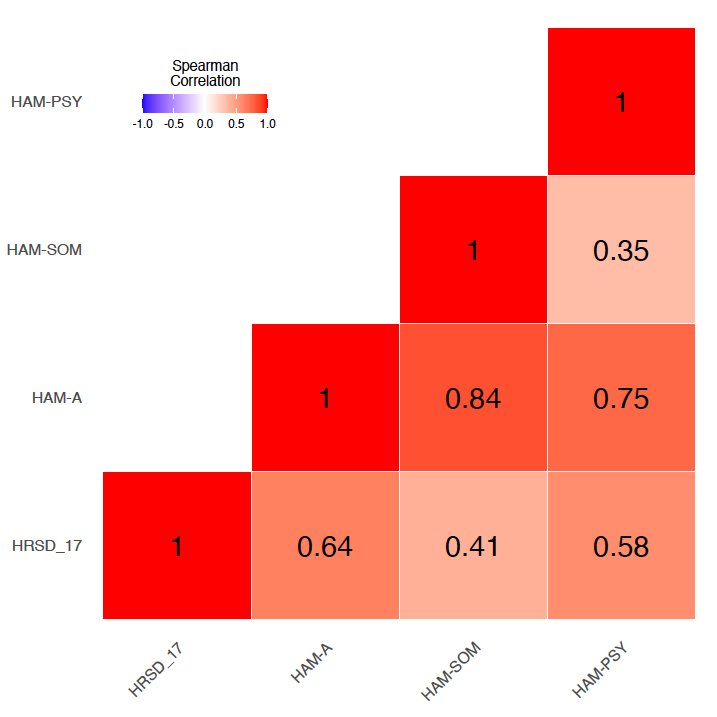
